# Testing hypothesized neural substrates with lesions

**DOI:** 10.64898/2026.11.18.767704

**Authors:** Patrik Bey, Joe Mole, James K. Ruffle, Amy Nelson, Edgar Chan, Lisa Cipolotti, Parashkev Nachev

## Abstract

Correlative neuroimaging generates anatomical hypotheses causally testable by lesion studies. Regionally weighted lesion volume-deficit regression—comparing slopes of behavioural dependence on damage inside vs outside a hypothesized neural substrate—and compartmental volume comparisons inheriting the same logic have been widely employed for this task, but without analytical examination or empirical validation of the underlying methodology. Here we show analytically that this approach does not provide a valid test of whether the hypothesized compartment identifies the true substrate. To illustrate the empirical consequences, we conduct semi-synthetic simulations using 4119 ischaemic lesions, modelling behavioural deficits dependent on a global lesion attribute, on widely investigated distributed cortical networks and white matter tracts, or on a right frontal substrate of fluid intelligence previously identified by multivariate methods. We apply regionally weighted regression with each hypothesized compartment as the target over 5000 iterations spanning multiple realistic noise levels. Where the hypothesized target is not the true substrate, we find spuriously significant dependence despite definitionally absent anatomical specificity; adjustment for lesion volume attenuates or exacerbates the observed bias dependent on the chosen target. Conversely, where the target is the true substrate, lesion volume correction universally reduces already poor sensitivity. Our analysis potentially explains common discrepancies across localisation studies, and highlights the necessity of multivariate approaches that explicitly model lesion and substrate anatomy at finely resolved spatial scales.

## 1. Introduction

No perfect method of functional brain mapping exists. Correlative methods, such as functional brain imaging, offer high anatomical precision but lack causal power; disruptive methods, such as lesion-deficit inference, can establish necessity but are constrained by the pathological context (Rorden & Karnath, 2004). One appealing solution is to assign each aspect of the inferential task to the method best suited to it, employing correlative data to *generate* hypotheses and disruptive data to *test* them.

A valid approach to such methodological synthesis is yet to be established, indeed none has been formally evaluated with the large-scale, semi-synthetic validation frameworks the problem requires (Mah et al., 2014; Pombo et al., 2023; Zavaglia et al., 2024). But many influential studies spanning several cognitive domains have adopted, with purely intuitive justification, a simple method of comparing the slopes of linear functions of the behaviour predicted by the extent of lesion overlap with vs outside the hypothesized substrate (e.g. (MacGregor et al., 2022; Reber et al., 2021; Roca et al., 2011; Smith et al., 2022; Woolgar et al., 2010, 2018)) or even simpler compartmental volume comparisons inheriting the same fundamental logic (e.g. (Biesbroek et al., 2013; Cassidy et al., 2018; Duering et al., 2011; Feng et al., 2015; Gajardo-Vidal et al., 2021; Griffis et al., 2017, 2017; Han et al., 2013; Ito et al., 2022; Lin et al., 2019; Marchina et al., 2011; Riley et al., 2011; Wang et al., 2013; Warren et al., 2014; Zhu et al., 2010)).

The validity of this “method of slopes”, more precisely regionally weighted lesion volume-deficit regression (RLDR), rests on the assumption that rejection of the null hypothesis of no difference between two regionally weighted lesion volume-deficit regression coefficients—one for a hypothesized target substrate, *H*, the second for the rest of the brain *∼H* (or another substrate, *H_2_*)— provides grounds for inferring the identity of the true substrate. But a radically different, untested substrate may yield identical or greater coefficient differences, and rejection of the null on the chosen *H* does not eliminate equal or better alternatives, indeed does not examine them at all. This is true of any comparison of prespecified coarse compartmental volumes.

Crucially, RLDR may yield a spuriously significant result *even when the true substrate is entirely outside the hypothesized regional compartment.* This is another manifestation of a cardinal aspect of lesion-deficit mapping, identified more than a decade ago (Mah et al., 2014), arising from the distributed structure of lesions and its interaction with the underlying neural substrates (Figure 1). Consider a lesion-deficit relation, for simplicity schematized in two dimensions, where injury to the true neural substrate, *T,* is accompanied by collateral damage to other regions of the brain, structured by the non-random anatomical patterns of the source pathology (Figure 1, left). Here mass-univariate analysis (voxel or prior region-of-interest-based) cannot distinguish the effects of damage to *T* from damage to collateral regions because it ignores correlations between voxels. It will therefore mistakenly infer a wholly irrelevant region, *A*, where the inherent structure of lesions incidentally lowers the variability of deficit-associated damage compared with the true substrate.

**Figure 1.**
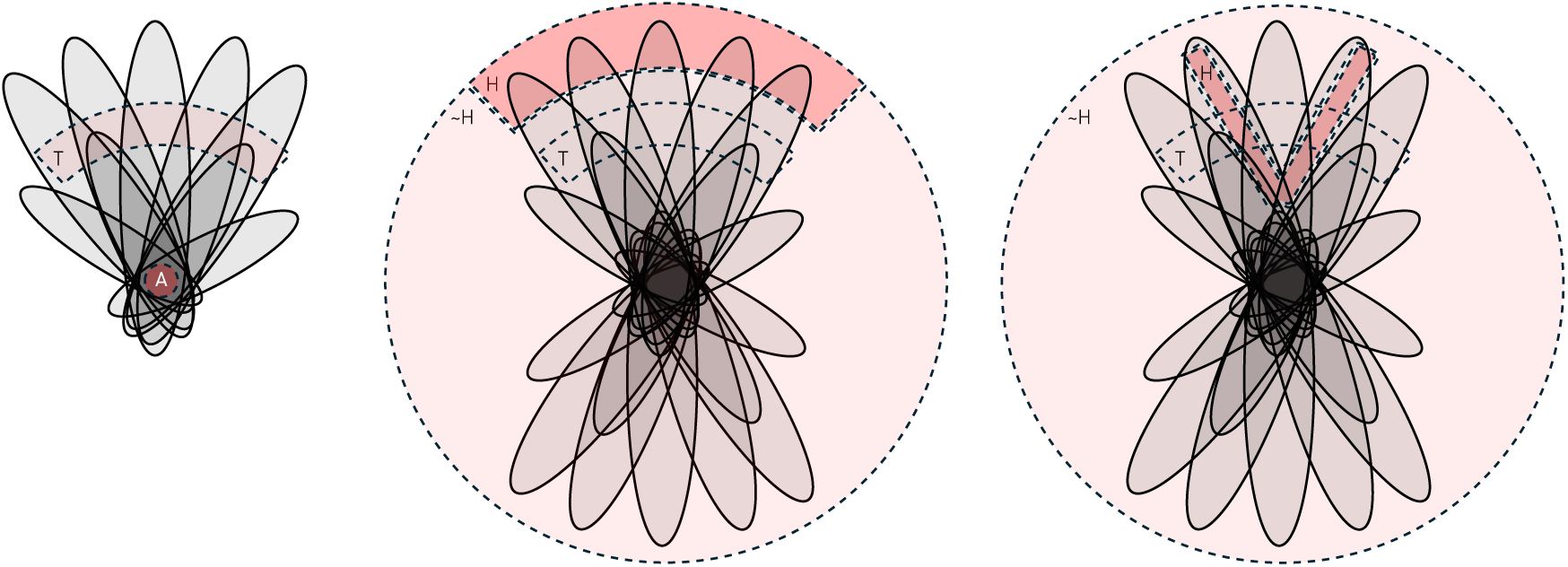
Illustration of the consequences of ignoring lesion and substrate topology in lesion-deficit inference. In voxel-wise mass-univariate lesion-deficit mapping (left), the pathology-driven distributed structure of lesions may cause dependence on the true substrate, *T*, to be mislocalised to a remote region, *A,* less variably correlated with the deficit (Mah et al., 2014). In regionally weighted lesion volume-deficit regression, a hypothesized substrate, *H,* may for the same reason be mistakenly inferred to be the true substrate, for both lesion and substrate topologies are assumed to respect the initial partitioning into *H* and *∼H*. This is true whether the substrate is a cortical network (middle) or a set of white matter tracts (right).

Compartmental volume comparisons not only ignore the topology of lesions, but presuppose that of the true substrate (Figure 1, middle and right) (Nachev, 2015). They project lesions into two pre-specified compartmental lesion volumes—hypothesized target and non-target—mistakenly assuming that a statistical comparison between their relations to the deficit is a valid test of target involvement. In fact, the difference in fitted slopes here only measures the difference in the strength of the relationship, so estimated, of each chosen compartment with the true, unknown substrate. Exceeding the threshold of statistical significance provides no warrant that the true substrate is within the target, let alone identical with it; equally, the test may be non-significant when the two perfectly coincide.

The foundational premise of lesion-deficit inference is that the behavioural deficit is generated by the interaction of two high-dimensional objects: the observed lesion and the (to be inferred) neural substrate. Any method that expresses this relation in terms of the lesioned volumes of *k* pre-specified compartments *compresses* both lesion and substrate to a *k*-dimensional representation. Where, as here, *k* = 2 (*H* vs *∼H*), this projection *must* lose behaviourally relevant information unless the true substrate consists of whole compartments, which is precisely what we are trying to infer and therefore cannot assume. The lost structure does not merely introduce imprecision but renders the anatomical relation of the true substrate to *H* unidentifiable.

This is formally provable under the minimal assumption that the deficit depends on the extent of substrate damage, with the same or varying sensitivity across the brain. For any set of lesions whose deficits arise from *H,* and where lesions involving *H* extend beyond it, there is another set of lesions with the same deficit scores and compartmental volumes under *H,* where the true substrate, *T*, lies outside *H*. This is because any multivoxel lesion that crosses the *H* vs *∼H* boundary can vary in its spatial configuration on either side of the boundary while retaining the same compartmental volumes, and therefore can damage the same volume of *T* as it did of *H* (or whatever volume differences in the substrate sensitivity of the deficit render equal in behavioural impact). Any method that sees only the *H* compartmental volumes and behavioural scores will therefore receive identical data and produce the same result irrespective of whether *H* or *T* obtains, whatever the model, volume covariates, threshold, or number of patients (Figure 2). Indeed, the lesions need not differ at all: Figure 2 shows the same lesions yielding the same compartmental volumes, and the same substrate damage, whether the substrate lies within *H* or wholly outside it. If this is true where *T* lies entirely outside *H*, it will be *a fortiori* true for a substrate that lies entirely within *H* yet is not identified by it, e.g. a small subset of a hypothesized distributed network.

**Figure 2.**
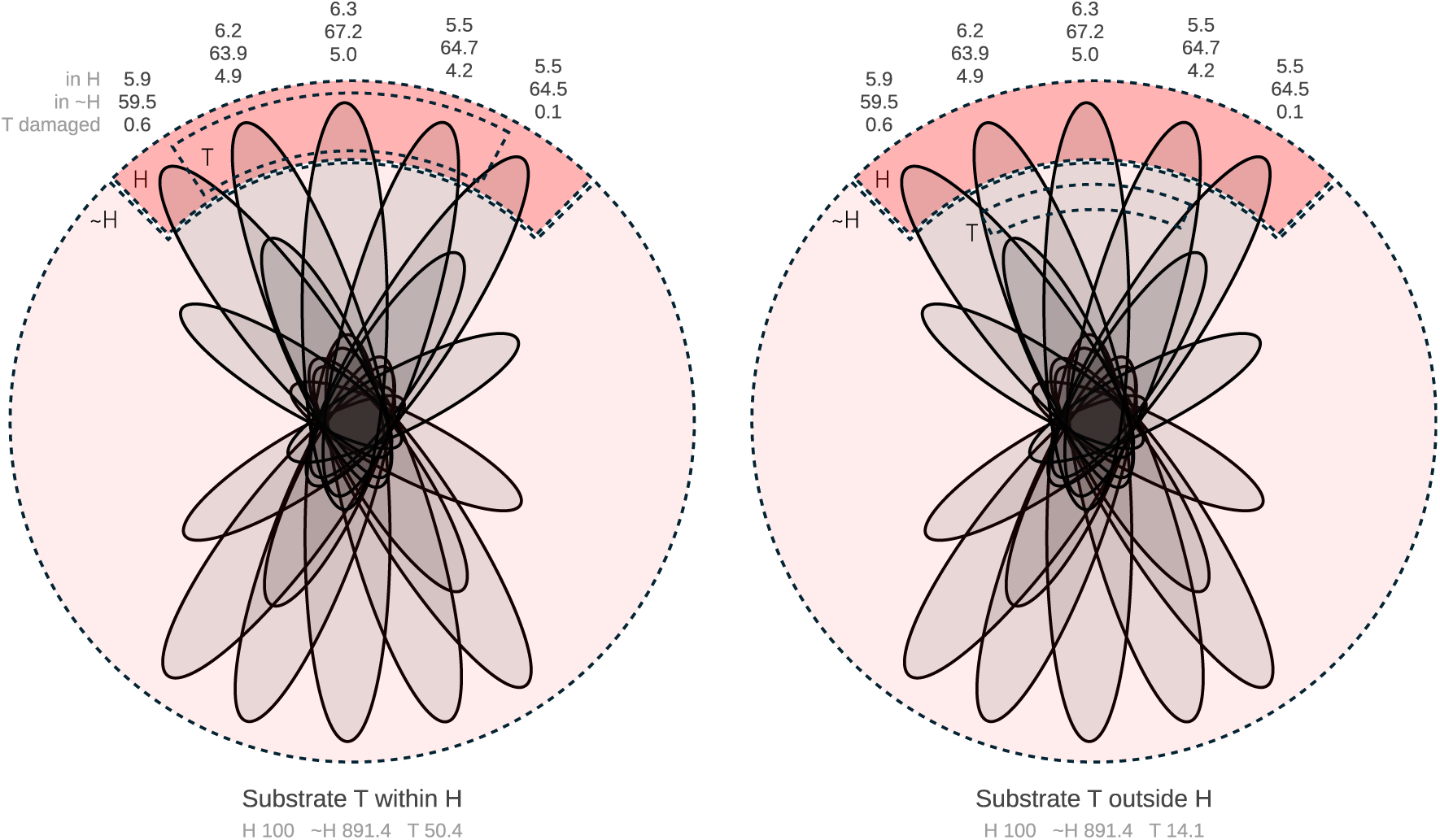
Illustration of how identical compartmental volumes may be obtained from radically different substrates. The schematized lesion-substrate relationship depicted in Figure 1 is replicated with two different true substrates, *T*, lying within *H* (left), or wholly outside it (right), with volumes in arbitrary units derived from the visualisation. Compartmental volumes with respect to *H* are identical across both scenarios, as is the volume of *T* damaged. A deficit determined by the volume of *T* damaged is therefore identical in every patient, irrespective of the location of *T*; so is one set by the proportional involvement, if tissue sensitivity is scaled by the ratio of the two substrates’ volumes (3.58), the factor by which the proportions differ for every lesion.

These criticisms are not hypothetical, in need of confirmation by subsequent empirical study, but analytic, inherent in the formal properties of the method no empirical evidence could change. The underlying defect has also been extensively investigated in other domains: this is the well-known “ecological fallacy” (Robinson, 1950) of attempting to infer from prespecified aggregate data (here hypothesized anatomical compartments) phenomena accessible only to individual analysis (here the finely resolved anatomy of lesions and substrates). The failure of identification it entails has long been established as irremediable through aggregate data alone: a definite answer rests on assumptions aggregate data cannot themselves verify (Cho & Manski, 2008; Cross & Manski, 2002; Goodman, 1953). The use of any *a priori* compartmentalisation presupposes functional homogeneity within compartments, precisely the anatomy we are supposed to infer (cf. (Elmalem et al., 2023)).

They also apply to any hypothesized substrate, any model of compartmental volumes, and any set of non-randomly distributed, spatially extended lesions: the only natural kind. But the risk of false positives will be higher the more widely distributed the hypothesized compartment and the more dissimilar its structure from the test lesion distribution: i.e. where multifocal *network* substrates or long white matter tracts are evaluated. Here target-intersecting lesions will typically be large and traverse multiple alternative substrates; conversely, non-target lesions will be highly heterogeneous in both volume and topology. False positives matter more than false negatives because rejecting the null hypothesis is (correctly) interpreted as more conclusive than failing to reject it (absence of evidence is not evidence of absence).

It is therefore instructive to demonstrate the consequences with semi-synthetic null models—involving real lesion data—of multifocal network substrates, where no regional lesion-deficit relation exists by design, and inference should yield no significant effect. We employ RLDR because simpler methods are confined to a subset of the compartmental information and therefore cannot escape criticism that arises from its poverty. We begin with the multiple demand network (MDN) as an illustrative example, owing to its scientific importance, and the role of RLDR as the only extant test of its necessity for behaviour. This large, distributed, bilateral, mainly frontal and parietal network is commonly activated by a variety of cognitively demanding tasks spanning perception, working memory and response selection. Following Luria (Luria, 1966), it is hypothesized to direct other regions in non-automatic tasks (Duncan, 2010; Duncan & Owen, 2000), to integrate cognitive components producing distinct executive functions (Assem et al., 2024), and to underpin one of the most important features of human cognition: fluid intelligence (Duncan et al., 2020; Duncan & Owen, 2000).

Applied to fluid intelligence, RLDR has provided support for the MDN as the critical substrate in a sample of 80 patients (Woolgar et al., 2010), but not in another sample of 227, where a hypothesis-free lesion-deficit mapping method implicated a predominantly right frontal network instead (Cipolotti et al., 2023). Differences in the data may conceivably account for the discrepancy, but further empirical replication alone would be inadequate, for methodological failure may produce consistent, even if erroneous, results where the defect, as here, lies in identification rather than in sample size (Mah et al., 2014; Xu, Jha, et al., 2018).

We conduct a large set of simulations, employing 4119 previously reported stroke lesions (Giles et al., 2025), that illustrate how RLDR spuriously implicates the MDN in the absence of true dependence on *any* functionally defined substrate. Where the MDN *is* the true substrate, we demonstrate that the method exhibits poor sensitivity, accentuated by covariate adjustment for lesion volume. We then examine how true dependence on a candidate substrate for fluid intelligence identified by a validated multivariate lesion-deficit method can yield a spurious association with the MDN, providing a potential explanation for discrepancies in the published literature. We proceed to illustrate the fundamental problem with an array of other commonly hypothesized networks and white matter tracts: the dorsal and ventral attention networks, the sensorimotor network, the superior and inferior longitudinal fasciculi, the arcuate fasciculus, and the inferior fronto-occipital fasciculus.

## 2. Materials and methods

Our objective is to illustrate the practical consequences of the analytically given defects we have identified. This requires an empirically informed, semi-synthetic generative model that combines real lesions with a realistic process of deriving an associated behavioural deficit under biologically plausible lesion-substrate relations. We cannot use real lesion-deficit pairs because we need to set a quantifiable ground truth against which the fidelity of the anatomical inference can be objectively assessed (Giles et al., 2025; Mah et al., 2014; Xu, Jha, et al., 2018; Zavaglia et al., 2024). We therefore proceed in four stages: assembly of a real lesion dataset; computation of morphological features and hypothesized target-substrate overlaps for each lesion; generation of semi-synthetic behavioural scores under three alternative generative models; and application of RLDR to 5000 independent Monte Carlo draws, under two model specifications (with and without lesion-volume covariate adjustment). Note only the behavioural score in the third stage is synthetic, derived from a specified model of the lesion-deficit relation. The analytical pipeline in the fourth stage reproduces the RLDR specification as reported in the studies that have used it. Full source code, including the exact MATLAB (R2025b), Python (3.13.11), and Bash (5.2.21) implementations of every step described below, is publicly available at https://github.com/high-dimensional/HypoSubstrateTesting.

### 2.1. Data

#### Lesions

We use a previously published dataset (Giles et al., 2025) of 4119 ischaemic stroke lesions, segmented from diffusion weighted imaging of a representative population with a radiologically confirmed acute ischaemic stroke (constrained only by MRI contraindication and tolerability), and non-linearly registered to Montreal Neurological Institute (MNI) space, yielding whole-brain, voxel-wise, binary, registered maps of ischaemic damage, resliced at 2 × 2 × 2 mm resolution. The lesion distribution is shown in Figure 3. Further details are provided elsewhere (Giles et al., 2025). Stroke is chosen as the commonest pathology in lesion-deficit studies.

**Figure 3.**
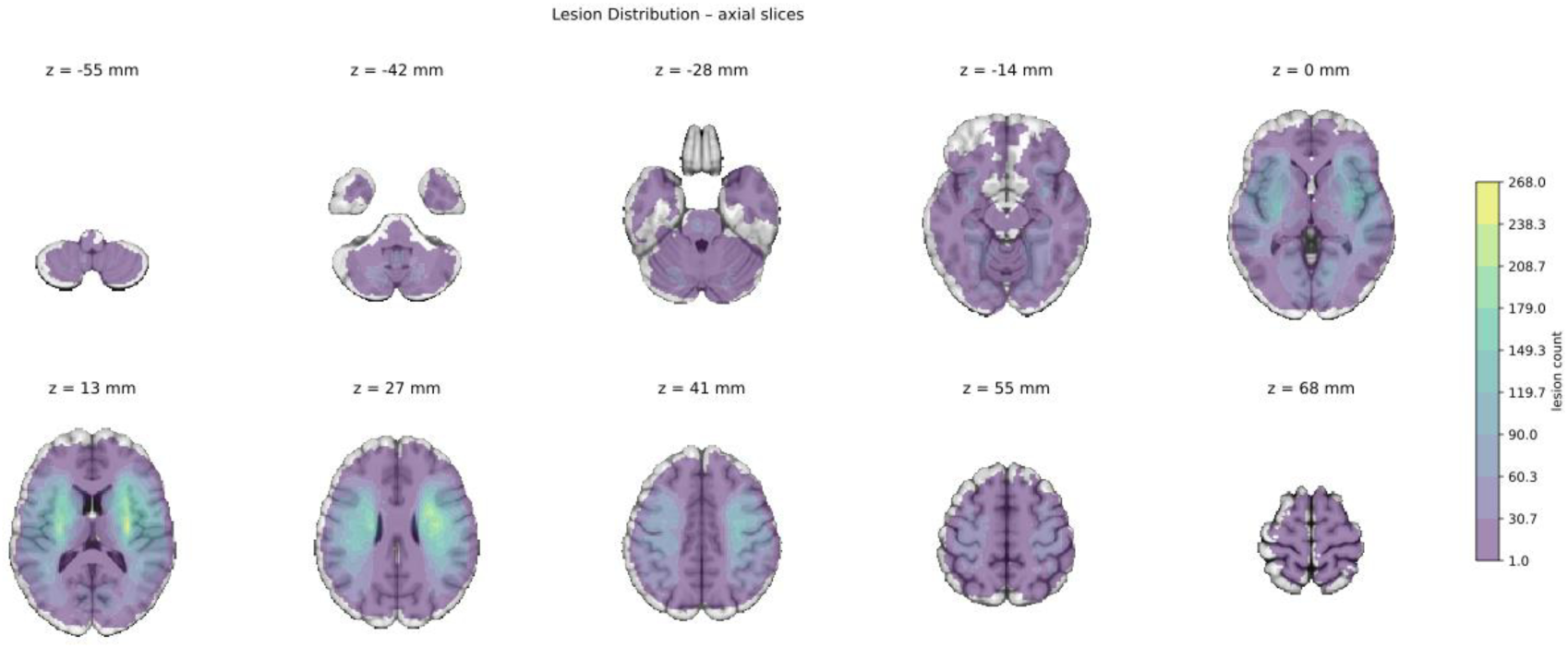
Lesion overlap map. The spatial distribution of the 4119 lesion masks used in the current cohort introduced in (Giles et al. 2025).

#### Substrate Maps

The multiple demand network (MDN) mask in MNI space was provided by Professor John Duncan’s research group and is the map used in (Woolgar et al., 2010). Masks of the dorsal attention, ventral attention, and sensorimotor networks were derived from the seven-network cortical parcellation of (Thomas Yeo et al., 2011). White-matter tract masks (superior longitudinal fasciculus, inferior longitudinal fasciculus, arcuate fasciculus, inferior fronto-occipital fasciculus) were derived from the Regionconnect atlas (Qi & Arfanakis, 2021). The right-frontal (RF) substrate mask is the network implicated in fluid intelligence by the multivariate graph lesion-deficit mapping reported in our previous study (Cipolotti et al., 2023).

### 2.2. Lesion features

For each of the 4119 lesions we compute two sets of features, both derived from the real binary lesion maps; these are the only inputs to the downstream simulations.

#### Morphological Features

MATLAB’s regionprops3 function was used to compute the largest principal axis length—a scalar measure of the lesion’s greatest spatial extent—as the lesion-size attribute used in the “global” generative scenario (§2.3). Where the lesion includes multiple connected components, the computed axis is of the ensemble taken as a whole.

#### Substrate-Overlap Features

For each lesion–hypothesized substrate pair we compute three scalar volumes with SPM25’s image calculator (spm_imcalc): the volume of overlap inside the hypothesized compartment, *v_in_* (voxels of the substrate mask damaged by the lesion); the volume of the lesion falling outside the substrate mask, *v_out_*; and their sum, the total lesion volume *v_tot_* = *v_in_* + *v_out_*. All three quantities are expressed in 10³ voxels. These are the compartmental lesion volumes on which RLDR is applied.

### 2.3. Semi-synthetic generative model

The derivation of the behavioural deficit score is the only synthetic element of the study, necessitated by the absence of any empirically available ground truth. This design preserves the realistic, pathology-driven, non-random distribution of lesions—the structure of which is the key source of inferential risk for RLDR—while allowing the ground-truth generator of behavioural deficit to be known by construction.

For each lesion *i*, a synthetic behavioural score *y_i_* is generated as

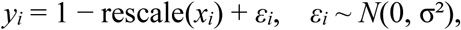

where rescale(·) performs min-max scaling of *x* across all 4119 lesions to the unit interval [0, 1], *ε_i_* is drawn independently from a zero-mean Gaussian distribution with per-iteration standard deviation σ, and *x_i_* is a scalar lesion feature whose definition distinguishes three generative scenarios. In the *global (lesion-size)* scenario, used in the analyses summarised by Figure 4, *x_i_* is the first principal axis length of lesion *i*; the score thus depends purely on lesion extent, with no regional specificity. In the *substrate-true* scenario, used in Figure 5 and in the right-hand (true-positive-rate) panels of Figure 7, *x_i_* is the volume of overlap between lesion *i* and the true substrate mask: the MDN for Figure 5, and each of the other cortical networks and white-matter tracts in turn for Figure 7. In the *RF-true* scenario, used in Figure 6, *x_i_* is the volume of overlap between lesion *i* and the right-frontal mask from Cipolotti et al. (2023).

**Figure 4.**
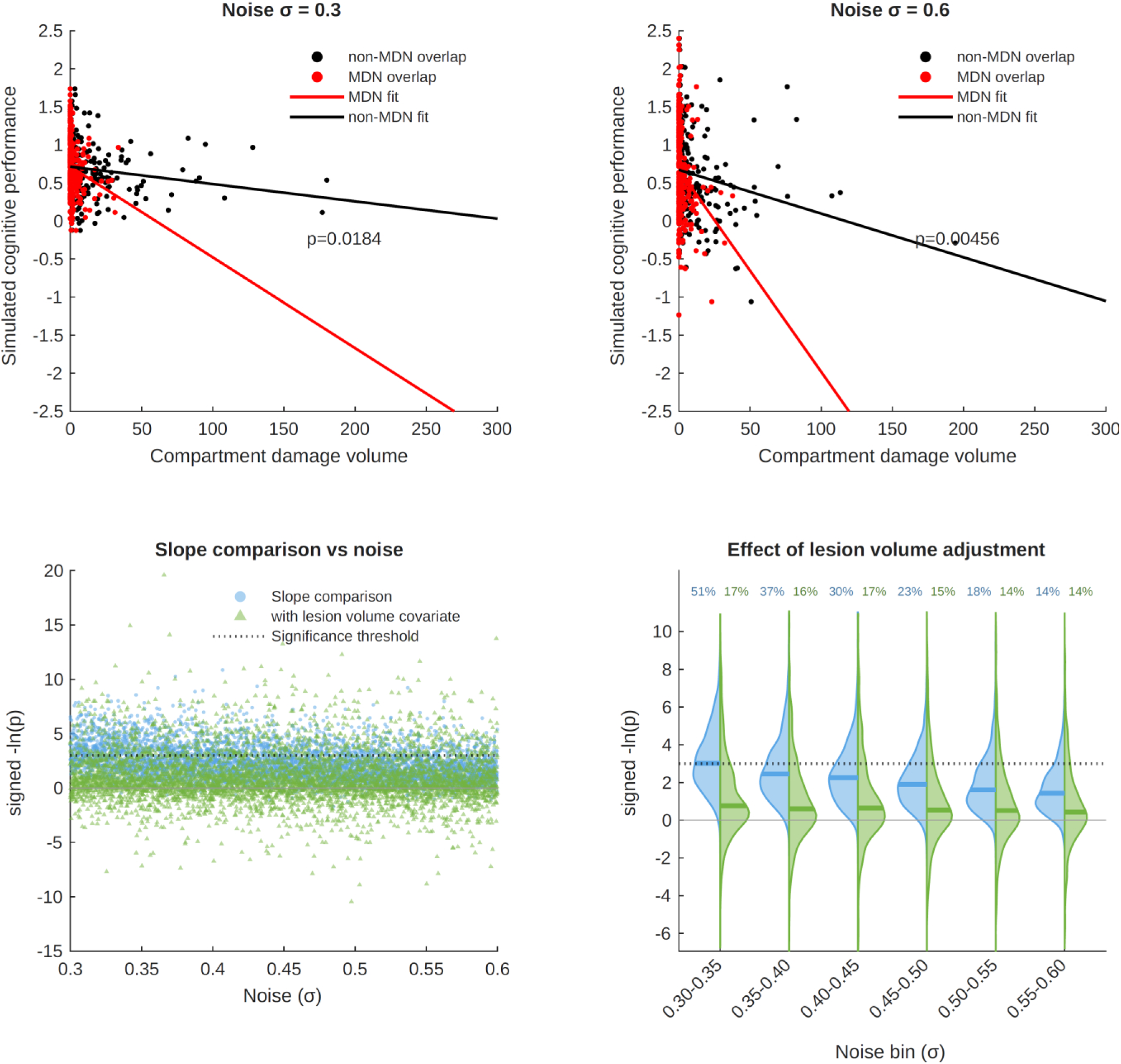
Semi-synthetic simulations of RLDR with MDN as the hypothesized substrate and lesion-size dependent deficit. Example RLDR simulations of a purely lesion-size dependent deficit at two different noise levels (σ=0.3, top left; σ=0.6, top right), showing linear fits to MDN (red) and non-MDN (black) compartmental volumes, and the *p* value for the difference between the slopes. The plots below show the *p* values (expressed as their negative log) from 5000 Monte Carlo simulations across the full noise range, with (green) and without (blue) the addition of lesion volume as a covariate in the model, plotted individually (bottom left), and binned as violin plots (bottom right). Note that the *p* value distributions exhibit substantial mass surviving the critical threshold of *p*<0.05 (dotted line), quantified in the binned plots separately for volume-adjusted and unadjusted simulations, resulting in frequent spurious inference of critical MDN involvement where there is none.

**Figure 5.**
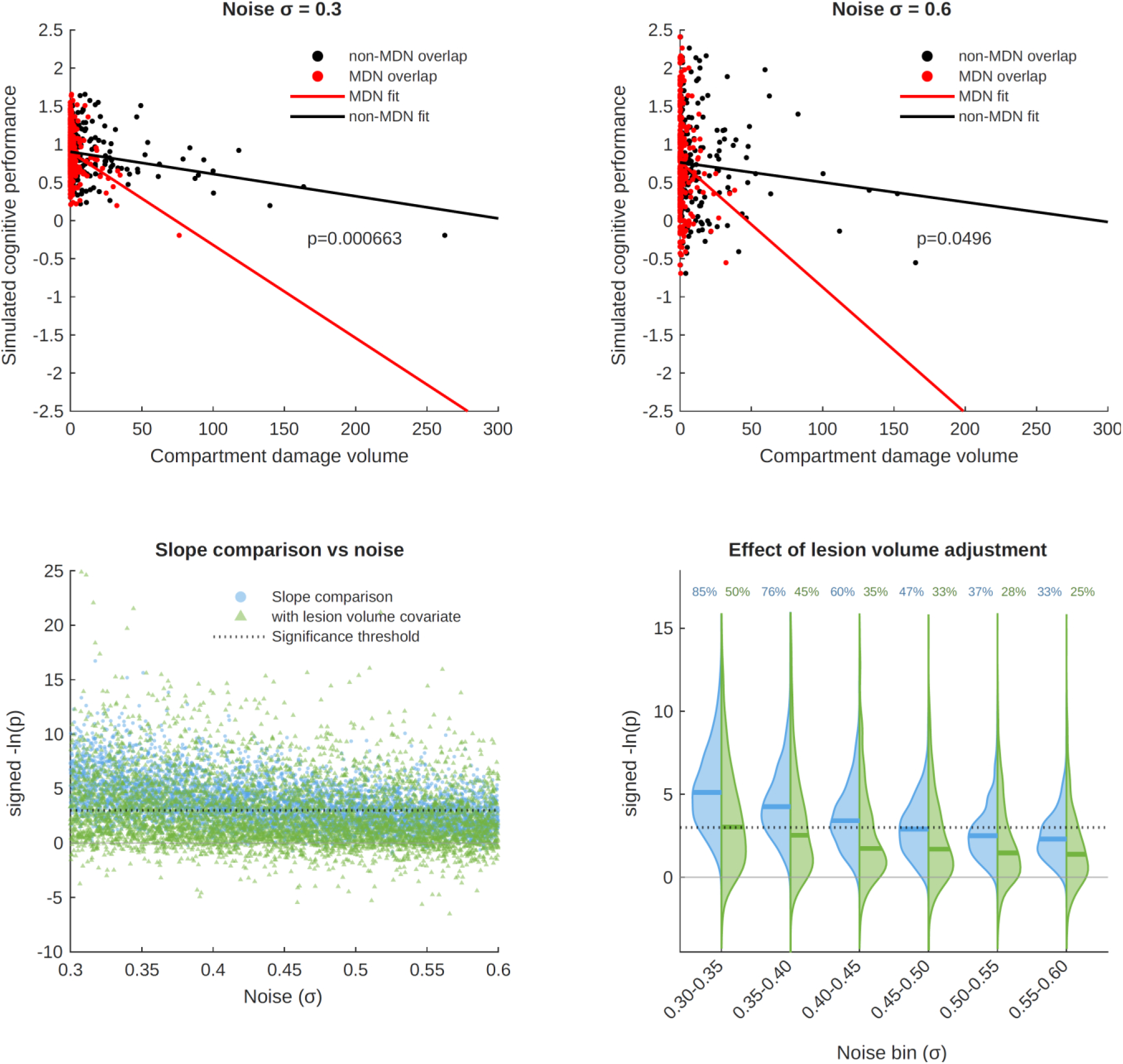
Semi-synthetic simulations of RLDR with MDN as the hypothesized and true substrate. Example RLDR simulations of an MDN-dependent deficit at two different noise levels (σ=0.3, top left; σ=0.6, top right), showing linear fits to MDN (red) and non-MDN (black) compartmental volumes, and the *p* value for the difference between the slopes. The plots below show the *p* values (expressed as their negative log) from 5000 Monte Carlo simulations across the full noise range, with (green) and without (blue) the addition of lesion volume as a covariate in the model, plotted individually (bottom left), and binned as violin plots (bottom right). Note that the *p* value distributions exhibit substantial mass failing to reach the critical threshold of *p*<0.05 (dotted line), quantified in the binned plots separately for volume-adjusted and unadjusted simulations, resulting in poor sensitivity for detecting true dependence, especially with adjustment for lesion volume.

**Figure 6.**
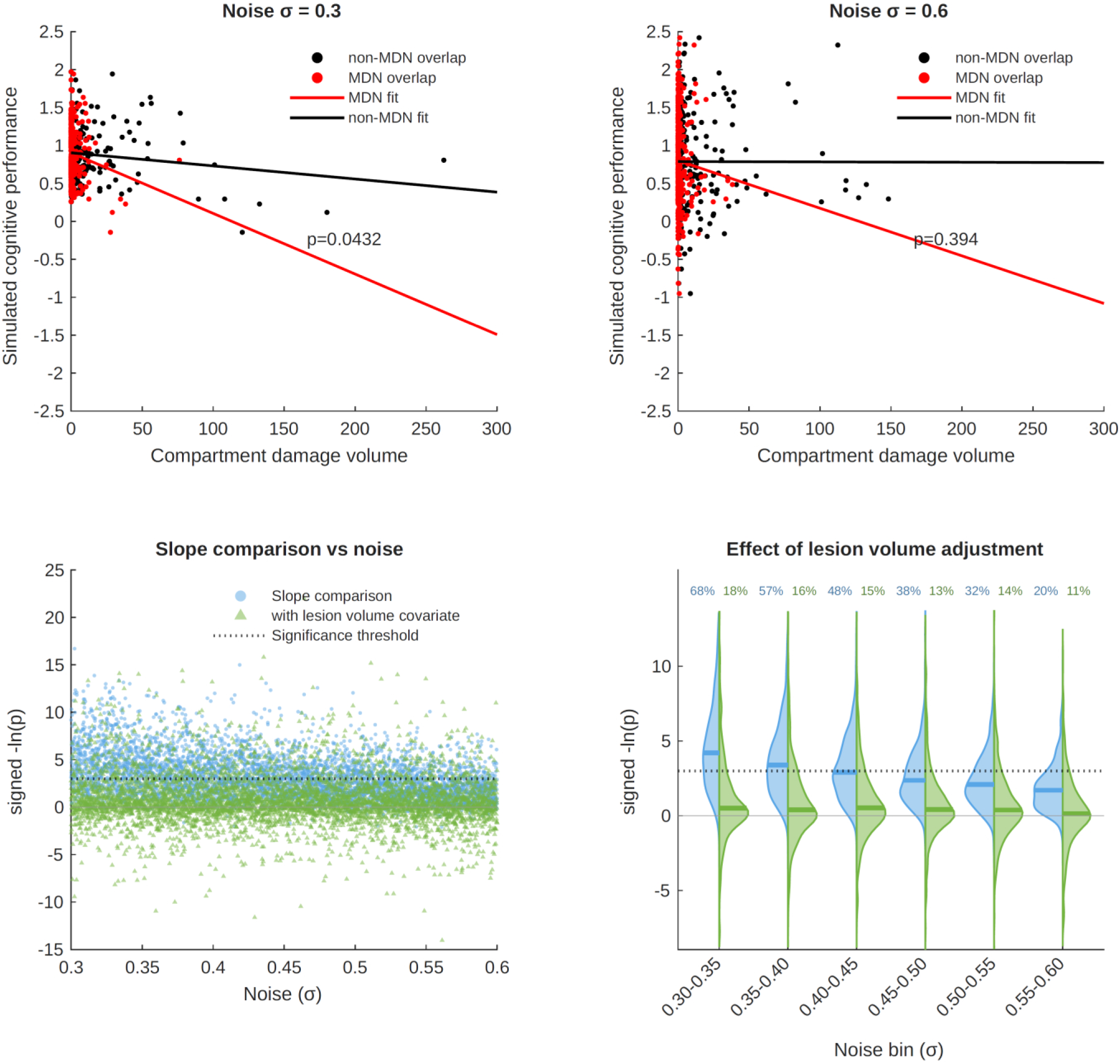
Semi-synthetic simulations of RLDR with MDN as the hypothesized substrate and a right frontal locus as the true. Example RLDR simulations of a right frontal substrate-dependent deficit at two different noise levels (σ=0.3, top left; σ=0.6, top right), showing linear fits to MDN (red) and non-MDN (black) compartmental volumes, and the *p* value for the difference between the slopes. The plots below show the *p* values (expressed as their negative log) from 5000 Monte Carlo simulations across the full noise range, with (green) and without (blue) the addition of lesion volume as a covariate in the model, plotted individually (bottom left), and binned as violin plots (bottom right). Note that the *p* value distributions exhibit substantial mass surviving the critical threshold of *p*<0.05 (dotted line), quantified in the binned plots separately for volume-adjusted and unadjusted simulations, resulting in frequent spurious inference of critical MDN involvement though the true substrate is an unrelated right frontal locus.

**Figure 7.**
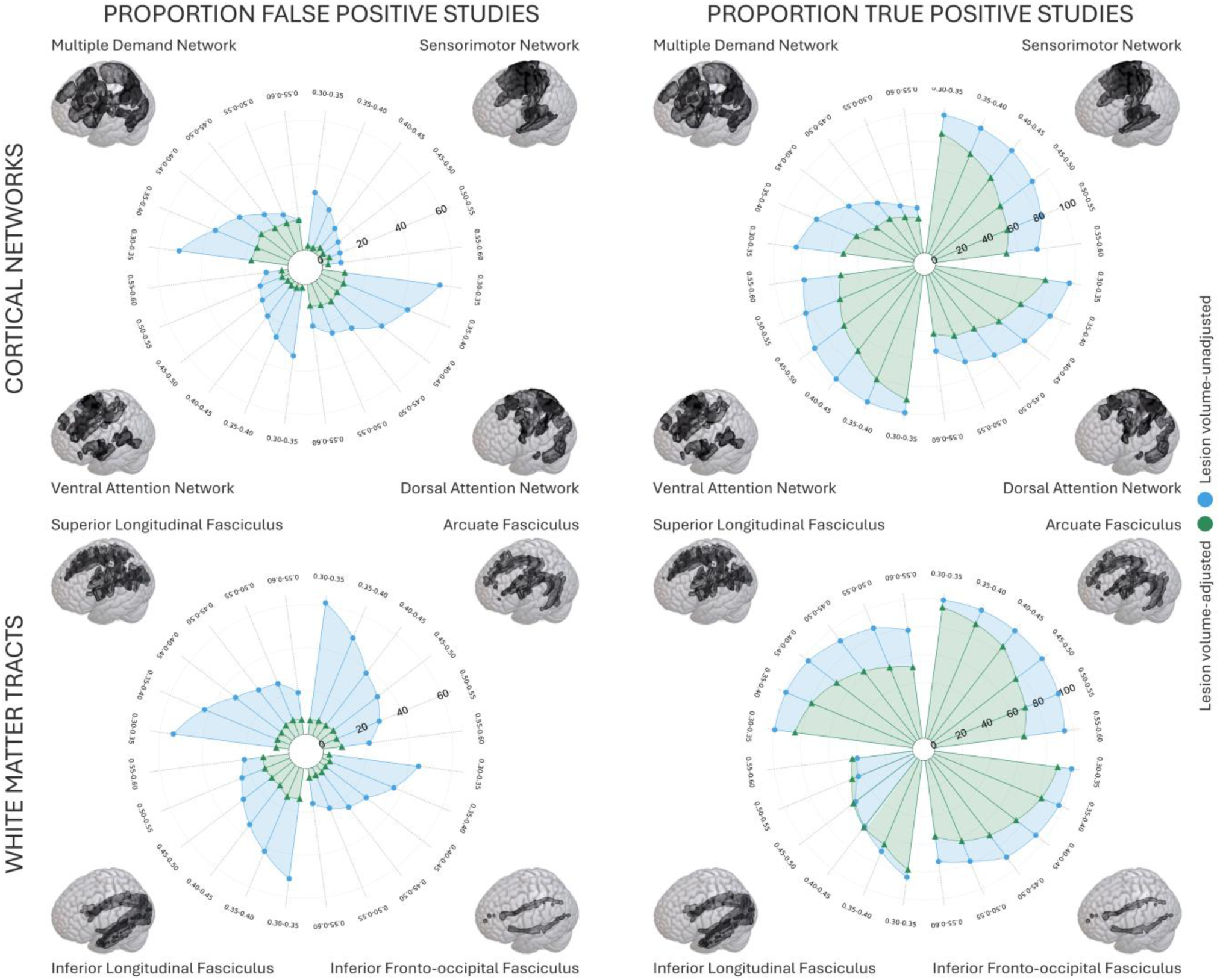
Generalisation across other networks, and white matter substrates. Simulations with parameters identical to the preceding (excluding the right frontal substrate ground truth) except for the hypothesized substrate—the dorsal and ventral attention networks, the sensorimotor network, the superior and inferior longitudinal fasciculi, the arcuate fasciculus, and the inferior fronto-occipital fasciculus—are summarized in radar lollipop plots, with the false positive rate (left) and true positive rate (right) as the radial dimension, and noise level in quadrants circumferentially. Note consistently high false positive rates and variable sensitivity for both cortical networks (top) and white matter tracts (bottom), and variable effects of adjustment for lesion volume. The MDN results are added for comparison.

Because *x_i_* is rescaled to [0, 1], the generative slope of *y_i_* on the rescaled feature is exactly −1 by construction in every scenario. We deliberately fix this slope rather than vary it: holding the effect size constant and varying only σ isolates the question of interest (how does RLDR behave as noise grows, for a fixed realistic ground truth?) from confounding variation in effect magnitude. A unit slope over a [0, 1] range is also a representative effect size in absolute terms, so that any failure of sensitivity (§3) reflects the method itself rather than an unrealistically under-powered generative model, and any false positive under an alternative truth (§3) reflects misattribution in the face of a strong alternative signal rather than floor-level noise. The range σ ∈ [0.3, 0.6] spans conditions from moderately noisy (σ = 0.3, such that the standard deviation of the noise is 30% of the full signal range) to heavily noisy (σ = 0.6, exceeding half the signal range), enclosing signal-to-noise regimes plausible for real behavioural measurements.

### 2.4. Monte Carlo simulations

We perform 5000 independent iterations per scenario. Iterations are indexed *j* = 1, …, 5000, with noise level σ*_j_* increasing linearly from 0.3 at *j* = 1 to 0.6 at *j* = 5000 (σ*_j_* = 0.3 + (*j* − 1) × 0.3 / 4999), so that the full noise range is sampled evenly across the run and results are resolved along the whole noise continuum.

Each iteration proceeds in three steps. First, a fresh noise vector *ε* is drawn for all 4119 lesions from *N*(0, σ*_j_*²), and synthetic scores *y_i_* are computed as described in §2.3. Second, to mimic a realistic lesion-deficit study design in which cohorts are enriched for impaired patients relative to the population we partition the 4119 lesions at the 30th percentile of the *noisy* scores *y_i_*, draw 150 lesions uniformly at random without replacement from the low-score group (*y_i_* ≤ the 30th percentile), and draw a further 150 from the high-score group (*y_i_* > the 30th percentile). The resulting 300-case sample balances impaired and unimpaired cases and is of a size typical of published lesion-deficit studies of fluid intelligence (e.g. *n* = 80 in Woolgar et al., 2010; *n* = 227 in Cipolotti et al., 2023). Third, two regression models are fitted to the 300-case sample, as described in §2.5, and the *p*-value of the target-vs-non-target slope-difference test is stored for each.

Iterations are independent: a fresh noise draw and a fresh sub-sample are generated at each step. A single global random seed (rng(42)) is fixed before the simulation loop, making the entire procedure deterministically reproducible.

### 2.5. RLDR evaluation

RLDR compares the slope of a linear regression of behavioural score on lesion–substrate overlap between the hypothesized target compartment and its complement. We reproduce this specification in its standard form, as applied in the literature we evaluate, by constructing a long-format data frame in which each patient contributes two observations (one per compartment), and fitting a single linear model with an interaction.

For each of the 300 sampled patients *i* we create a target row, with Overlap = *v*_in,i_ (overlap with the target compartment), Substrate = 1 (target indicator), Volume = *v*_tot,i_ (the patient’s total lesion volume), and Behaviour = *y*_i_; and a non-target row, with Overlap = *v*_out,i_ (overlap outside the target), Substrate = 0 (non-target indicator), Volume = *v*_tot,i_ (the same total lesion volume), and Behaviour = *y*_i_ (the same simulated score). On the resulting 600-row table we fit two models using MATLAB’s fitlm with the “interactions” specification. Writing *O*_ic_, *S*_ic_ and *V*_i_ for the Overlap, Substrate and Volume of patient *i*’s row for compartment *c* ∈ {target, non-target}, the unadjusted model (Model 1) regresses Behaviour on Overlap, Substrate, and their interaction:

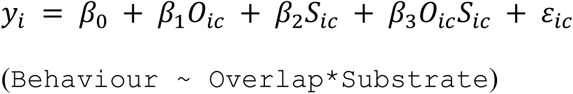

The volume-adjusted model (Model 2) additionally includes Volume and its interactions with Overlap and Substrate as nuisance terms, reflecting the practice of adjusting for total lesion volume:

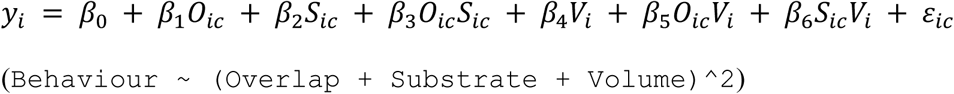

In both, the errors *ε*_ic_ are taken as independent and identically distributed, as fitlm assumes.

In both models, the coefficient *β*_3_ on the Overlap × Substrate interaction tests whether the regression slope of Behaviour on Overlap differs between the target and non-target compartments: the slope is *β*_1_ + *β*_3_ for the target and *β*_1_ for the non-target in Model 1, and *β*_1_ + *β*_3_ + *β*_5_*V* and *β*_1_ + *β*_5_*V* at any given volume *V* in Model 2. This is the slope-difference test on which RLDR rests. We extract its p-value from a Type III ANOVA of the fitted model in MATLAB; for this highest-order, single-degree-of-freedom term it coincides with the two-sided t-test of *β*_3_. For a directional hypothesis—the expected kind here—the p-value is halved, yielding a one-sided test: a result counts as positive only where the target slope is significantly the steeper of the two slopes. A significant difference in the opposite direction is treated as a negative, for the complement *∼H* is not itself a hypothesis about the target substrate. We apply the standard *p* < 0.05 threshold and store the statistic −ln(*p*) for downstream summarisation. No multiple-comparison correction is applied, reflecting the real-world use of the method in which a single pre-specified test is run per study. Exactly one p-value per model, per iteration, per scenario is therefore produced.

### 2.6. Hypothesis testing scenarios

We evaluate seventeen hypothesis testing scenarios, each comprising 5000 iterations fitted under both Model 1 and Model 2 (yielding 10,000 fits per condition). Three primary scenarios use the MDN as the hypothesized target. The first, MDN-hypothesized with the global (lesion-size) generator (Figure 4), tests the false-positive rate under a lesion-size-driven deficit with no anatomical specificity. The second, MDN-hypothesized with the MDN as the true generator (Figure 5), tests the true-positive rate when the hypothesis is correct. The third, MDN-hypothesized with the RF locus as the true generator (Figure 6), tests the false-positive rate under a realistic alternative substrate drawn from prior multivariate work on the same behavioural construct. Seven further conditions extend the analysis to other candidate substrates, cortical or white matter-based: the dorsal attention, ventral attention, and sensorimotor networks, and the superior longitudinal, inferior longitudinal, arcuate, and inferior fronto-occipital fasciculi. Each of these is evaluated twice, once with the global-true generator and once with the same substrate as the true generator, yielding fourteen further runs summarised jointly with the MDN conditions in Figure 7.

## 3. Results

Where the deficit is driven solely by a global feature of the lesion, RLDR may show a statistically significant difference *without* any specific dependence on MDN anatomy, under both low- (Figure 4, top left) and high- (Figure 4, top right) noise conditions. Across 5000 simulations spanning a wide range of noise levels, spurious statistical differences commonly arise (Figure 4, bottom), and are not adequately controlled by covariate adjustment with another global lesion parameter such as lesion volume. Much of the density of the *p* value distribution (ranging 51% to 14% across noise levels) survives the critical statistical threshold in the unadjusted case, and a substantial proportion of it (ranging from 17% to 14% across noise levels) in the adjusted case. A plausible, wholly non-regional confounding effect here tends to result in erroneous inference of MDN substrate dependence.

Where the MDN is modelled as the true substrate, 85% to 33%, across noise levels, of simulated studies yield a significant association, dropping to 50% to 25% with lesion volume adjustment (Figure 5). This illustrates the distortion adjusting for lesion volume inevitably introduces owing to the complex relationship between lesion topology and volume (Xu, Jha, et al., 2018). Even when the MDN is the true substrate, RLDR cannot be relied upon to detect it.

Where a deficit is driven by another candidate substrate—the right frontal network we have previously identified (Cipolotti et al., 2023)—RLDR may erroneously infer MDN involvement (Figure 6, top). Across the 5000 simulations spurious statistical differences frequently arise at every noise level (Figure 6, bottom), mirroring the effects observed in the global scenario. The proportion of spurious associations is substantial for both volume unadjusted (68-20%) and adjusted (18-11%) models. MDN dependence can be mistakenly inferred where the ground truth substrate is a right frontal locus, not the MDN.

This pattern is not unique to the MDN. Across both other cortical networks and white matter substrates, high false positive rates are commonly observed, sometimes ameliorated and sometimes exacerbated by lesion volume adjustment (Figure 7). Equally, where the target is the true substrate, sensitivity is frequently low, and universally impaired by lesion volume adjustment. Individual substrate plots are provided in Supplementary Figures S1-S14.

## 4. Discussion

Lesion-deficit mapping is a challenging task. Operating under a tight data regime, it involves inference of unknown, likely complex functional topologies, from sets of noisy lesion-deficit pairs generated by partially observed, obviously complex pathological processes. However understandable, the temptation to look for simplifying shortcuts must be resisted, for the fundamental nature of the system prohibits them. The distributed structure of lesions and neural substrates cannot be assumed to be simple, as mass-univariate lesion-deficit mapping does explicitly, and compartmental volume comparisons do implicitly. Misinference is the likely result, as rigorous methodological evaluations invariably show (Mah et al., 2014; Pombo et al., 2023; Zavaglia et al., 2024).

Any compartmental method, RLDR included, presupposes the spatial organisation it attempts to infer. As a matter of basic topology, neither the significance nor the direction of any statistical comparison is a reliable guide to the location of the true substrate, regardless of statistical model choice, threshold selection, covariate adjustment, or number of patients. If the same compartmental scores— definitionally the sole input of the method—can be produced by radically different true substrates, neither positive nor negative results are interpretable, and nothing following the compartmentalisation step could conceivably remedy the defect.

This conclusion is analytic, rendered inevitable by the core process of *a priori* aggregation, whose consequences have been recognized in the broader inferential literature for three quarters of a century. It also mirrors familiar analytic criticisms of mass-univariate and lesion volume-adjusted lesion-deficit inference. The former assumes lesion covariance is either unstructured or Gaussian, an *a priori* imposed structure of a different kind but with analogous inferential effects (Mah et al., 2014; Nachev, 2015; Xu, Jha, et al., 2018); the latter is effectively a global compartmental parameter and introduces formally the same error (Husain & Nachev, 2007; Mah et al., 2014; Xu, Jha, et al., 2018).

Although our focus is the testing of compartmentally defined functional hypotheses, the criticism extends to any method based on prespecified regions of interest, in proportion to the potential substrate diversity the size of the parcellation admits. Increasing the resolution of the parcellation, to the asymptote of single voxels, transfers the inferential burden to the (inevitably multivariate) model of regional interactions. Where the parcellation is not prespecified but derived from a multivariate model of the lesion and substrate anatomy informed by the behaviour of interest (e.g. (Cipolotti et al., 2023)), compartmental homogeneity is inferred rather than assumed and the concern does not arise.

We have seen that the defect of prespecified compartmental comparisons is irremediable because it arises in the very first, foundational step. No modification to the statistical model can correct an error caused by loss of information at the input. Correcting for total lesion volume, or any other low dimensional lesion parameter, invokes a crudely aggregated attribute raising the same objection, and— as our semi-synthetic simulations illustrate—cannot provide an effective remedy. In any event, adjustment for lesion volume inevitably penalises topologies correlated with it, replacing one source of distortion with a potentially greater other (Husain & Nachev, 2007; Mah et al., 2014; Xu, Jha, et al., 2018). We see this effect vividly in the impact on sensitivity where the target is the true substrate.

We have also seen that the defect is one of identifiability, not statistical efficiency. Neither larger cohorts, nor replication over multiple studies, could increase our confidence in the result, whether negative or positive. Nor is this a matter of the precision with which lesions are segmented, compartments defined, or individual anatomies registered, for the compression compartmentalisation induces is far more lossy than any preceding step. Equally, although every lesion-deficit model is improved by better behavioural characterisation, the problem here lies entirely on the anatomical side. Even where the hypothesized region is highly correlated with the behaviour, all we can conclude is that the true substrate is commensurately correlated, not that it has been identified: lesion and substrate topologies remain entangled.

A special case arises where the hypothesized compartments are large enough to eliminate lesions lying across them, e.g. hemispheric compartments with unilateral lesions, or crude antero-posterior divisions with lesions at sublobar scale. If we set connectivity aside, we can reasonably conclude that the true substrate lies somewhere within the compartment. Few, however, would accept that conditional, even if they accepted the very low anatomical precision so crude a compartmentalisation imposes. Assembling cohorts with small lesions respecting the hypothesized boundary is a luxury the underlying pathology rarely affords.

It may reasonably be objected that aggregation is unavoidable given the spatially extended nature of lesions themselves, at least as conventionally modelled in the form of binary anatomical masks. But that the properties of natural lesions impose a limit on the spatial resolution with which a neural substrate can be inferred does not license us to aggregate the substrate itself *a priori*. Lesions are moreover not simple blobs, but exhibit a complex structure learnable with the right multivariate model and data of sufficient scale and coverage (Bonkhoff et al., 2021; Cipolotti et al., 2023; Giles et al., 2025; Pombo et al., 2023). How closely we can approach the substrate resolution limit depends on the fidelity of that model, and such prior (typically very weak) as we can credibly impose on the substrate itself. This is the core of the argument for multivariate models in lesion-deficit mapping we have elaborated elsewhere at length (Mah et al., 2014; Nachev, 2015; Pombo et al., 2023; Xu, Jha, et al., 2018).

Given a multivariate lesion-deficit model, the question is what form *a priori* hypothesis testing could take. Since such models yield an inferred map directly, the hypothesis can be evaluated against the map itself rather than through a compartmental contrast. Where the architecture admits an explicit prior, one could compare models with and without the hypothesis in the prior, quantified by model evidence or out-of-sample prediction. In any event, the comparison is informative only to the extent the map is identifiable from the lesions: where alternative maps fit equally well, the evidence for a hypothesis reflects the concentration of the prior rather than the data themselves. A hypothesis should therefore be compared with alternatives of matched size and complexity.

To identify fatal flaws in RLDR is not to conclude that the MDN is not the critical substrate of the abilities attributed to it. Since lesions provide the *only* source of causal evidence here, the loss is nonetheless important, indeed arguably deprives the theory of its most rigorous test. The same is true of the other substrates we examine.

In short, questions such as “does the degree of involvement of substrate X predict behaviour Y” cannot be reliably answered by a simple summary comparison of the properties of lesions differentiated by their overlap with X. Expressive, multivariate lesion-deficit models are necessary, whose integration with correlative data remains the object of future research. Given the immense complexity of the underlying problem—topological inference in the setting of richly structured, noisy, biased, incomplete data—no simple solution is plausible, and any candidate approach will inevitably require extensive numerical investigation within the semi-synthetic frameworks now established as the only robust means of validating lesion-deficit mapping of the human brain (Mah et al., 2014; Pombo et al., 2023; Xu, Rolf Jäger, et al., 2018).

## Acknowledgements

We are grateful to Professor John Duncan for supplying the MDN map used in the study.

