## Supplementary Material for "Testing hypothesized neural substrates with lesions"

Patrik Bey[1], Joe Mole[1,2], James K. Ruffle[1,3], Amy Nelson[1], Edgar Chan[1,2], Lisa Cipolotti,[1,2] and Parashkev Nachev[1]

Author affiliations:

1 Department of Translational Neuroscience and Stroke, Queen Square Institute of Neurology, WC1N 3BG, London, United Kingdom

2 Department of Neuropsychology, National Hospital for Neurology and Neurosurgery, WC1N 3BG, London, United Kingdom

3 Lysholm Department of Neuroradiology, National Hospital for Neurology and Neurosurgery, WC1N 3BG, London, United Kingdom.

Correspondence to: Parashkev Nachev or Lisa Cipolotti

Department of Translational Neuroscience and Stroke, Queen Square Institute of Neurology, WC1N 3BG, London, United Kingdom

#### 1. Lesion morphology associated semi-synthetic simulations

Semi-synthetic simulations of behaviour driven purely by a global lesion parameter across various cortical networks and white matter tracts as described in the main text (see Figure 4).

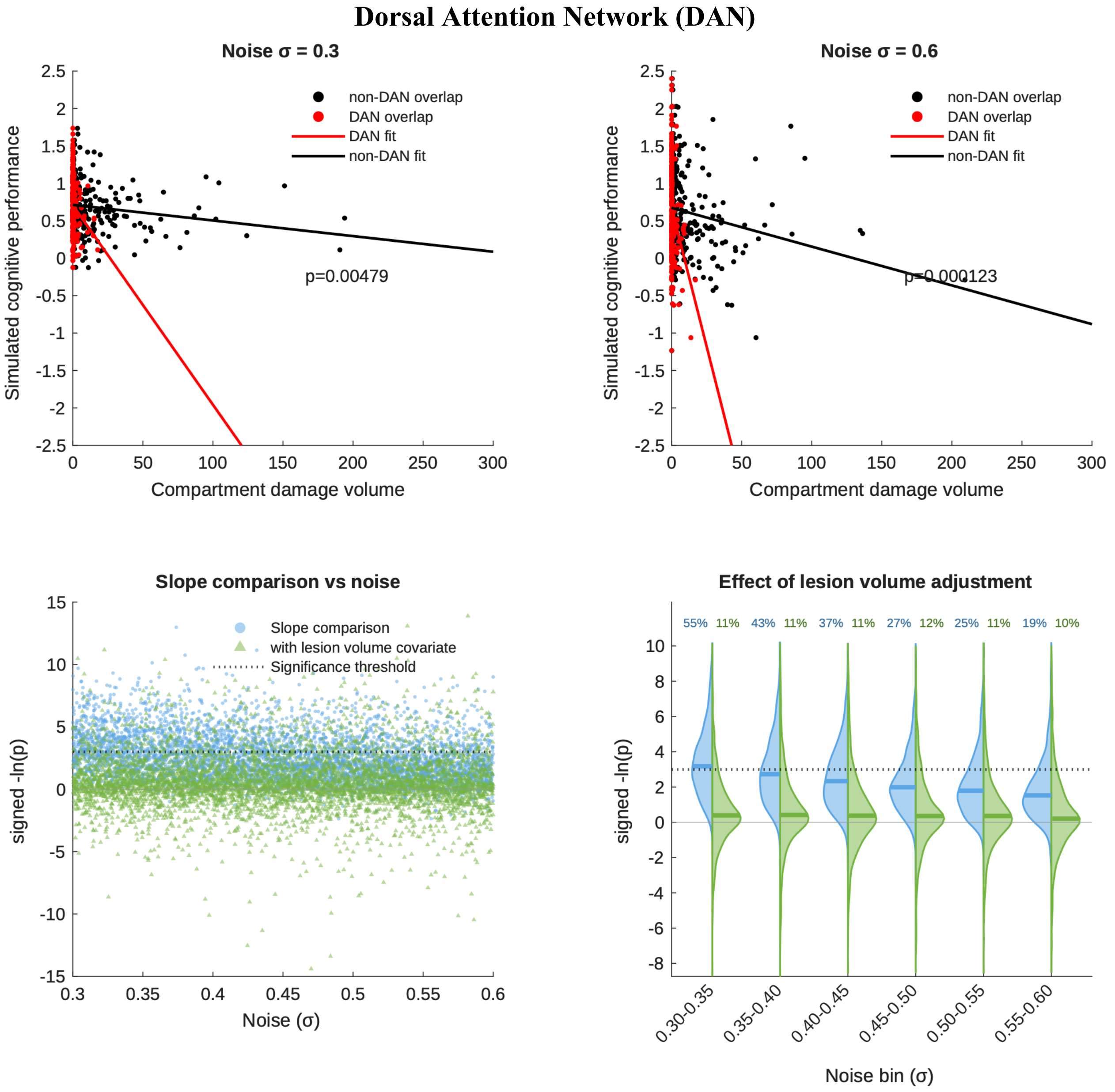


**Figure S1. Semi-synthetic simulations of RLDR with the Dorsal Attention Network (DAN) as the hypothesized substrate and lesion-size dependent deficit.** Example RLDR simulations of a purely lesion-size dependent deficit at two different noise levels (σ=0.3, top left; σ=0.6, top right), showing linear fits to DAN (red) and non-DAN (black) compartmental volumes, and the p value for the difference between the slopes. The plots below show the p values (expressed as their negative log) from 5000 Monte Carlo simulations across the full noise range, with (green) and without (blue) the addition of lesion volume as a covariate in the model, plotted individually (bottom left), and binned as violin plots (bottom right). Note that the p value distributions exhibit substantial mass surviving the critical threshold of $p<0.05$ (dotted line), quantified in the binned plots separately for volume-adjusted and unadjusted simulations, resulting in frequent spurious inference of critical DAN involvement where there is none.

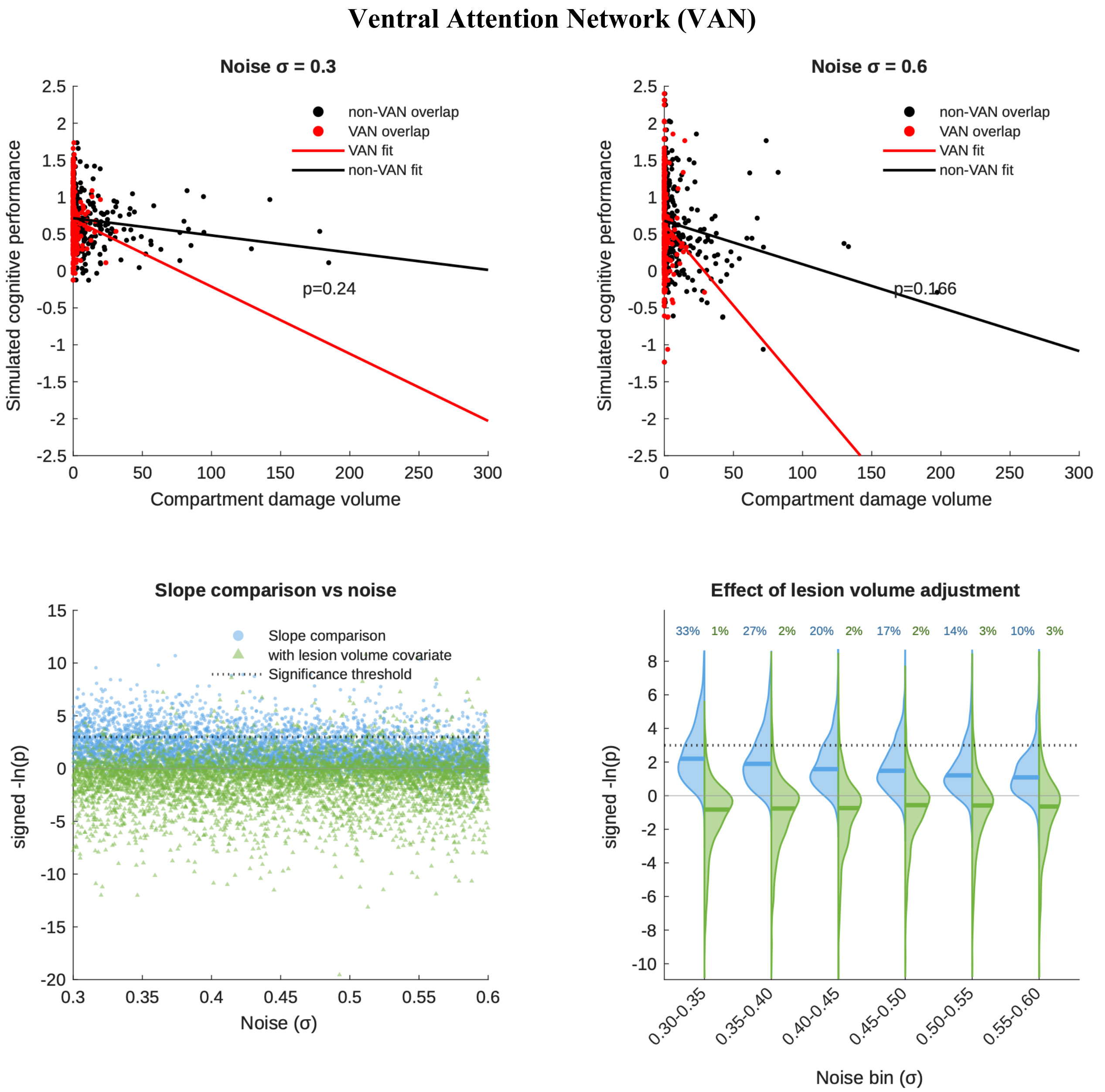


**Figure S2. Semi-synthetic simulations of RLDR with the Ventral Attention Network (VAN) as the hypothesized substrate and lesion-size dependent deficit.** Example RLDR simulations of a purely lesion-size dependent deficit at two different noise levels (σ=0.3, top left; σ=0.6, top right), showing linear fits to VAN (red) and non-VAN (black) compartmental volumes, and the p value for the difference between the slopes. The plots below show the p values (expressed as their negative log) from 5000 Monte Carlo simulations across the full noise range, with (green) and without (blue) the addition of lesion volume as a covariate in the model, plotted individually (bottom left), and binned as violin plots (bottom right). Note that the p value distributions exhibit substantial mass surviving the critical threshold of $p<0.05$ (dotted line), quantified in the binned plots separately for volume-adjusted and unadjusted simulations, resulting in frequent spurious inference of critical VAN involvement where there is none.

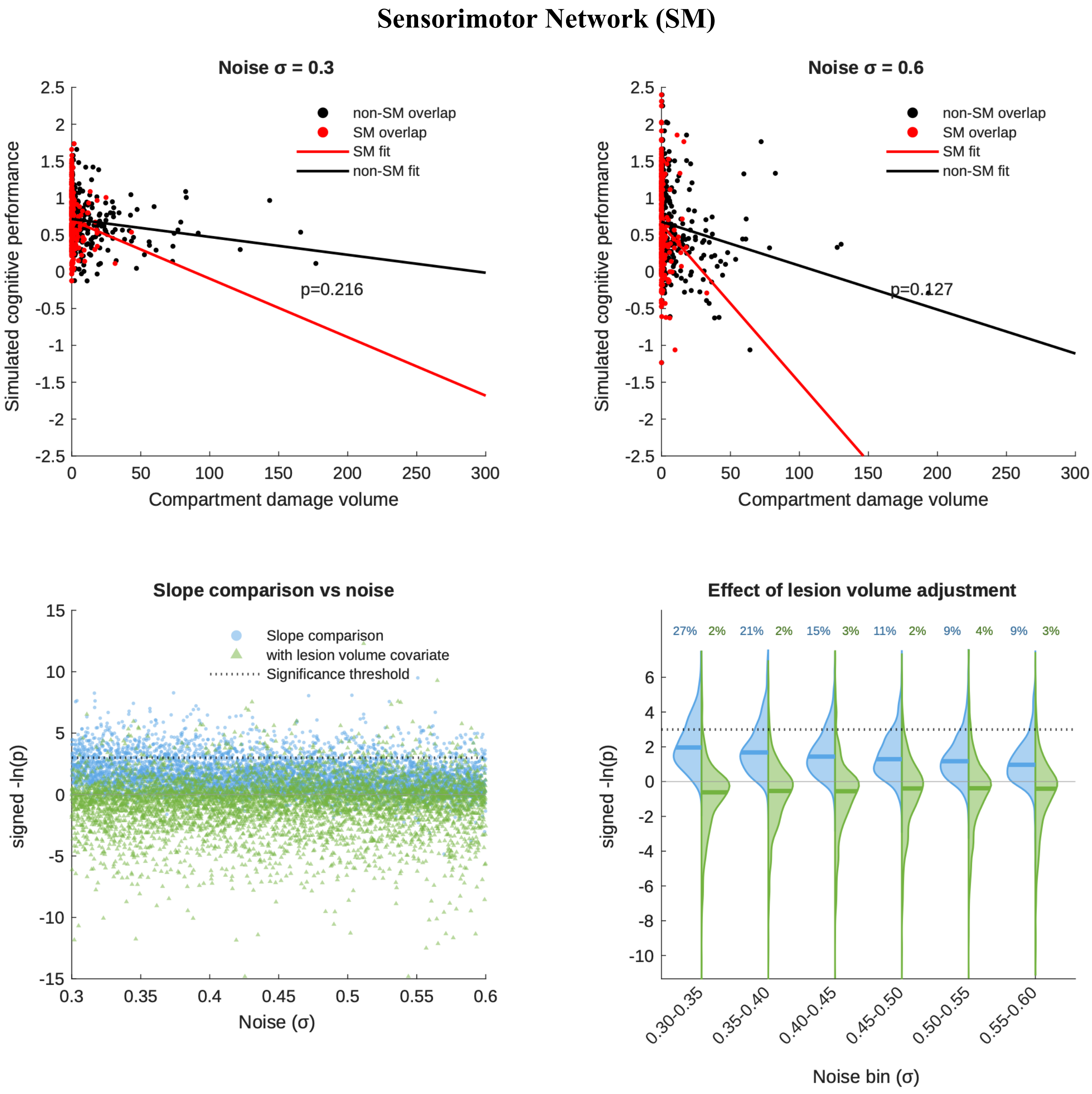


**Figure S3. Semi-synthetic simulations of RLDR with the Sensorimotor Network (SM) as the hypothesized substrate and lesion-size dependent deficit.** Example RLDR simulations of a purely lesion-size dependent deficit at two different noise levels (σ=0.3, top left; σ=0.6, top right), showing linear fits to SM (red) and non-SM (black) compartmental volumes, and the p value for the difference between the slopes. The plots below show the p values (expressed as their negative log) from 5000 Monte Carlo simulations across the full noise range, with (green) and without (blue) the addition of lesion volume as a covariate in the model, plotted individually (bottom left), and binned as violin plots (bottom right). Note that the p value distributions exhibit substantial mass surviving the critical threshold of $p<0.05$ (dotted line), quantified in the binned plots separately for volume-adjusted and unadjusted simulations, resulting in frequent spurious inference of critical SM involvement where there is none.

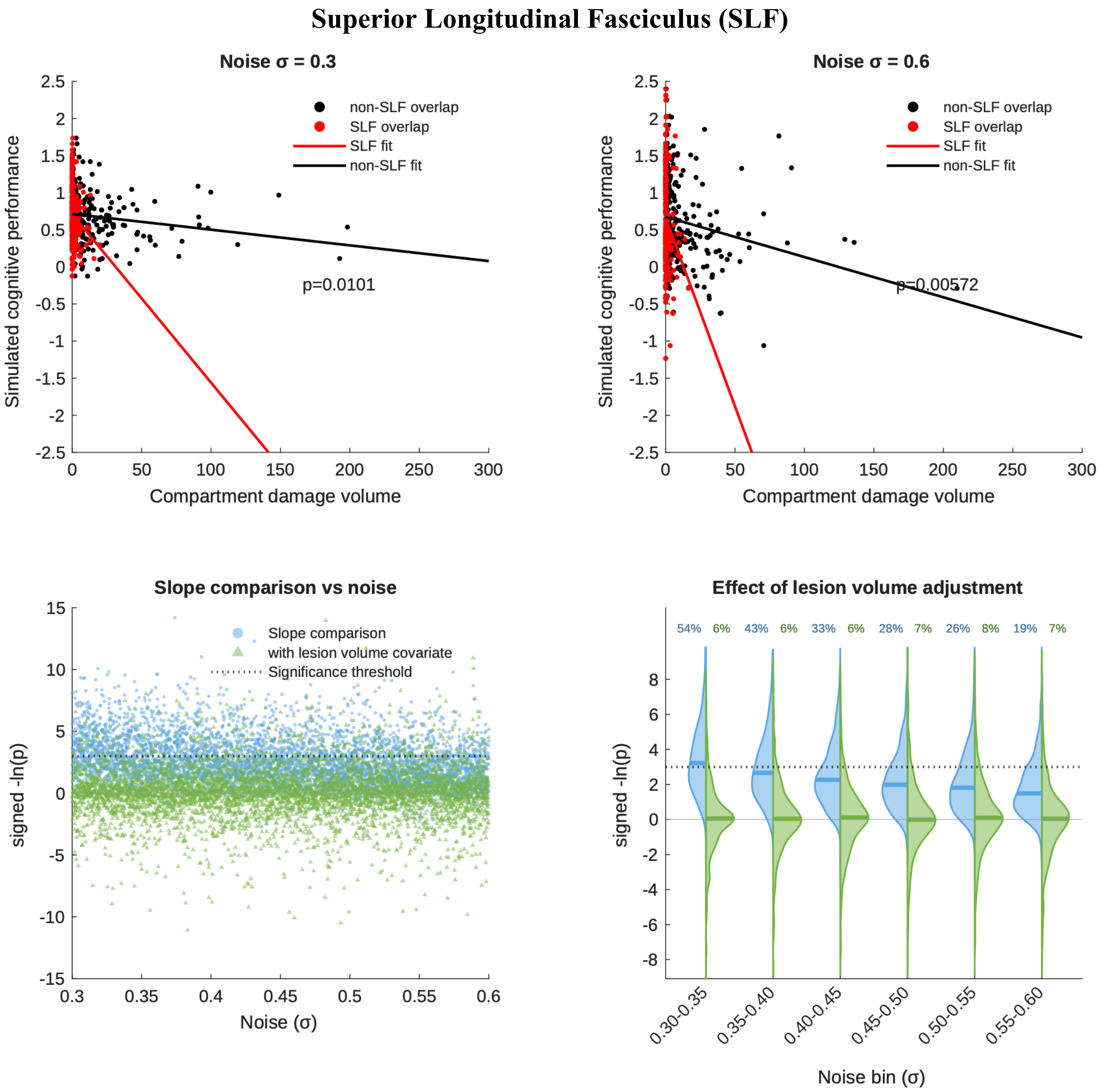


**Figure S4. Semi-synthetic simulations of RLDR with the Superior Longitudinal Fasciculus (SLF) as the hypothesized substrate and lesion-size dependent deficit.** Example RLDR simulations of a purely lesion-size dependent deficit at two different noise levels (σ=0.3, top left; σ=0.6, top right), showing linear fits to SLF (red) and non-SLF (black) compartmental volumes, and the p value for the difference between the slopes. The plots below show the p values (expressed as their negative log) from 5000 Monte Carlo simulations across the full noise range, with (green) and without (blue) the addition of lesion volume as a covariate in the model, plotted individually (bottom left), and binned as violin plots (bottom right). Note that the p value distributions exhibit substantial mass surviving the critical threshold of $p<0.05$ (dotted line), quantified in the binned plots separately for volume-adjusted and unadjusted simulations, resulting in frequent spurious inference of critical SLF involvement where there is none.

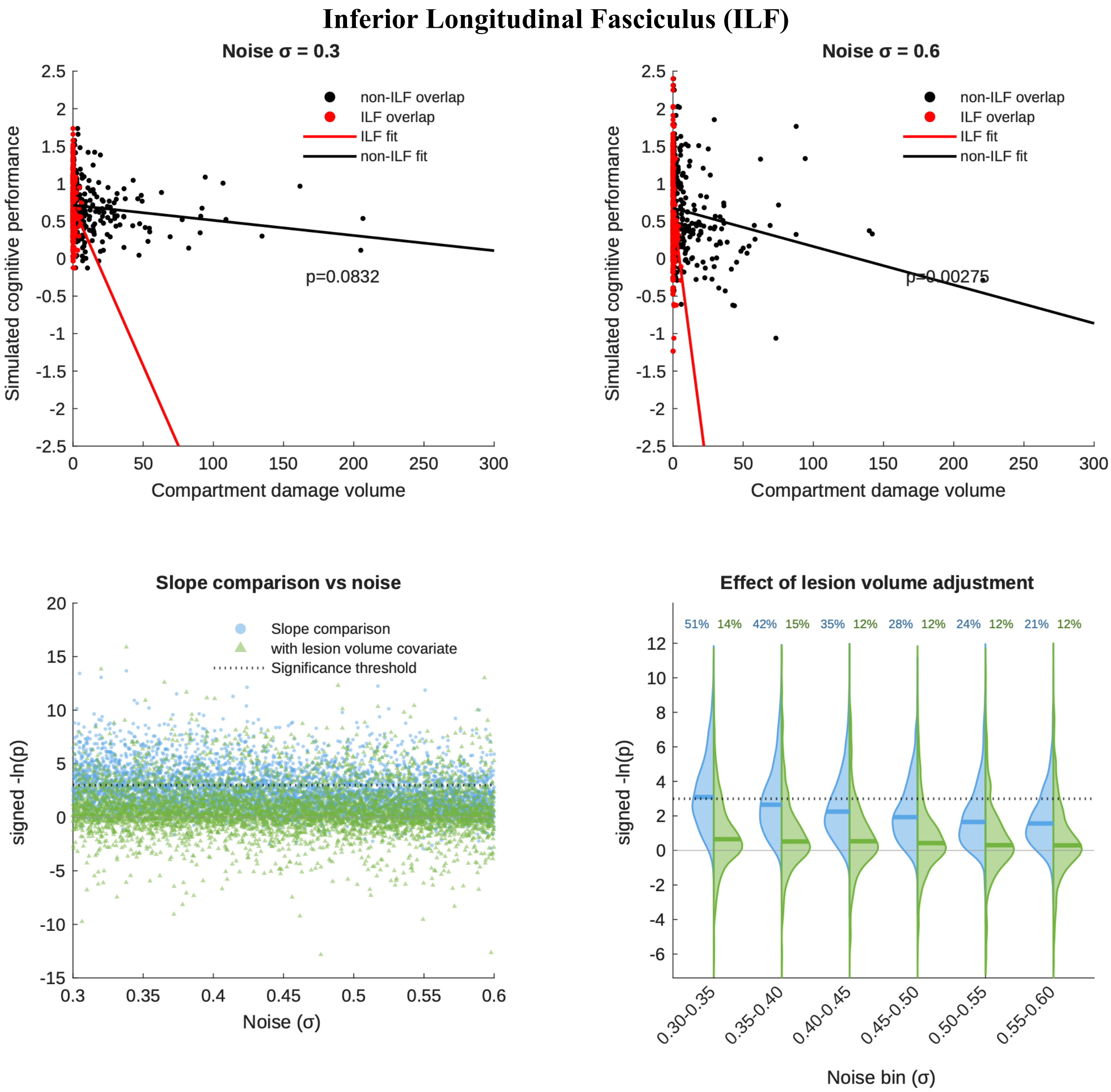


**Figure S5. Semi-synthetic simulations of RLDR with the Inferior Longitudinal Fasciculus (ILF) as the hypothesized substrate and lesion-size dependent deficit.** Example RLDR simulations of a purely lesion-size dependent deficit at two different noise levels (σ=0.3, top left; σ=0.6, top right), showing linear fits to ILF (red) and non-ILF (black) compartmental volumes, and the p value for the difference between the slopes. The plots below show the p values (expressed as their negative log) from 5000 Monte Carlo simulations across the full noise range, with (green) and without (blue) the addition of lesion volume as a covariate in the model, plotted individually (bottom left), and binned as violin plots (bottom right). Note that the p value distributions exhibit substantial mass surviving the critical threshold of $p<0.05$ (dotted line), quantified in the binned plots separately for volume-adjusted and unadjusted simulations, resulting in frequent spurious inference of critical ILF involvement where there is none.

### Arcuate Fasciculus (AF)

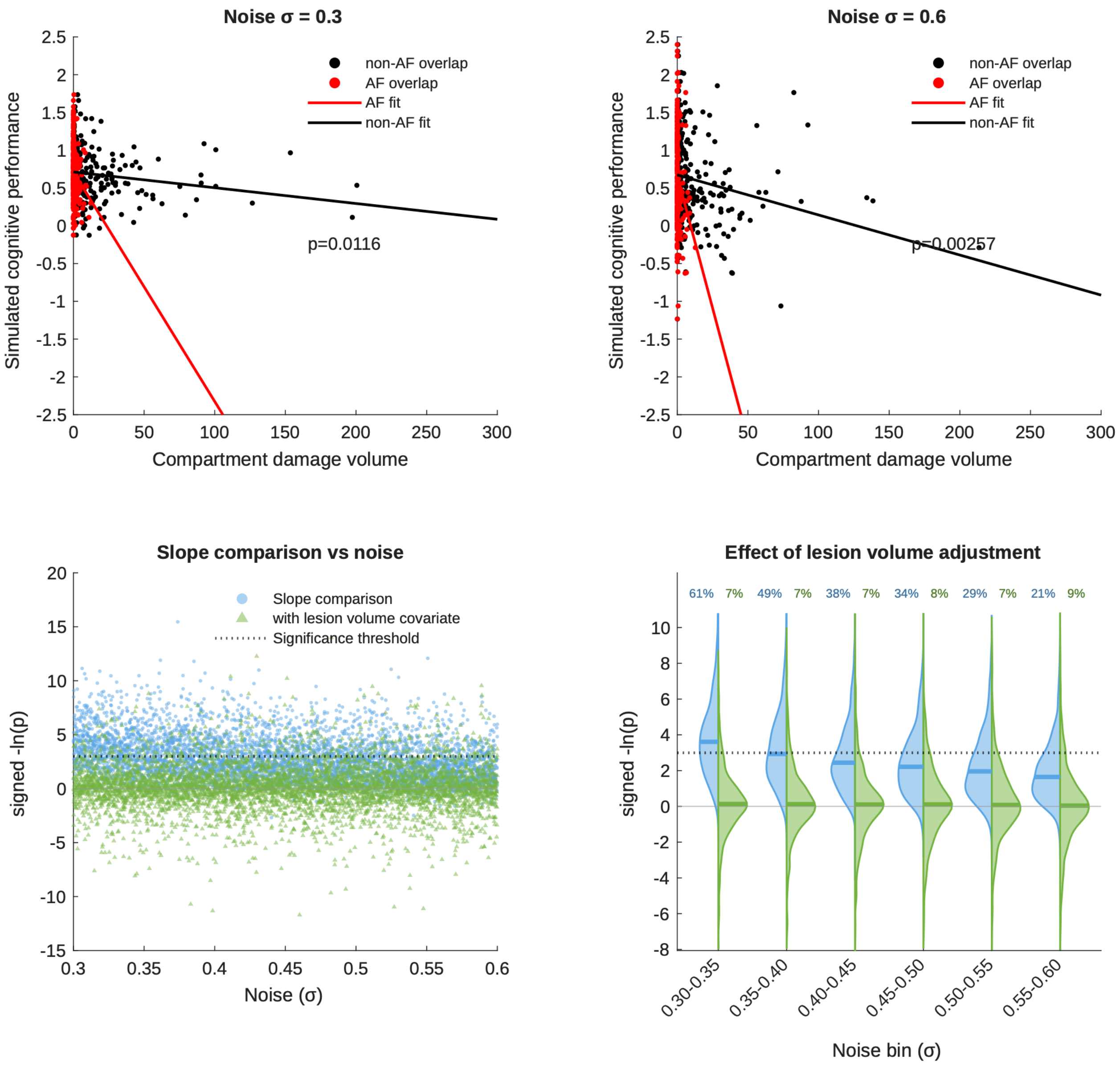


**Figure S6. Semi-synthetic simulations of RLDR with the Arcuate Fasciculus (AF) as the hypothesized substrate and lesion-size dependent deficit.** Example RLDR simulations of a purely lesion-size dependent deficit at two different noise levels (σ=0.3, top left; σ=0.6, top right), showing linear fits to AF (red) and non-AF (black) compartmental volumes, and the p value for the difference between the slopes. The plots below show the p values (expressed as their negative log) from 5000 Monte Carlo simulations across the full noise range, with (green) and without (blue) the addition of lesion volume as a covariate in the model, plotted individually (bottom left), and binned as violin plots (bottom right). Note that the p value distributions exhibit substantial mass surviving the critical threshold of $p<0.05$ (dotted line), quantified in the binned plots separately for volume-adjusted and unadjusted simulations, resulting in frequent spurious inference of critical AF involvement where there is none.

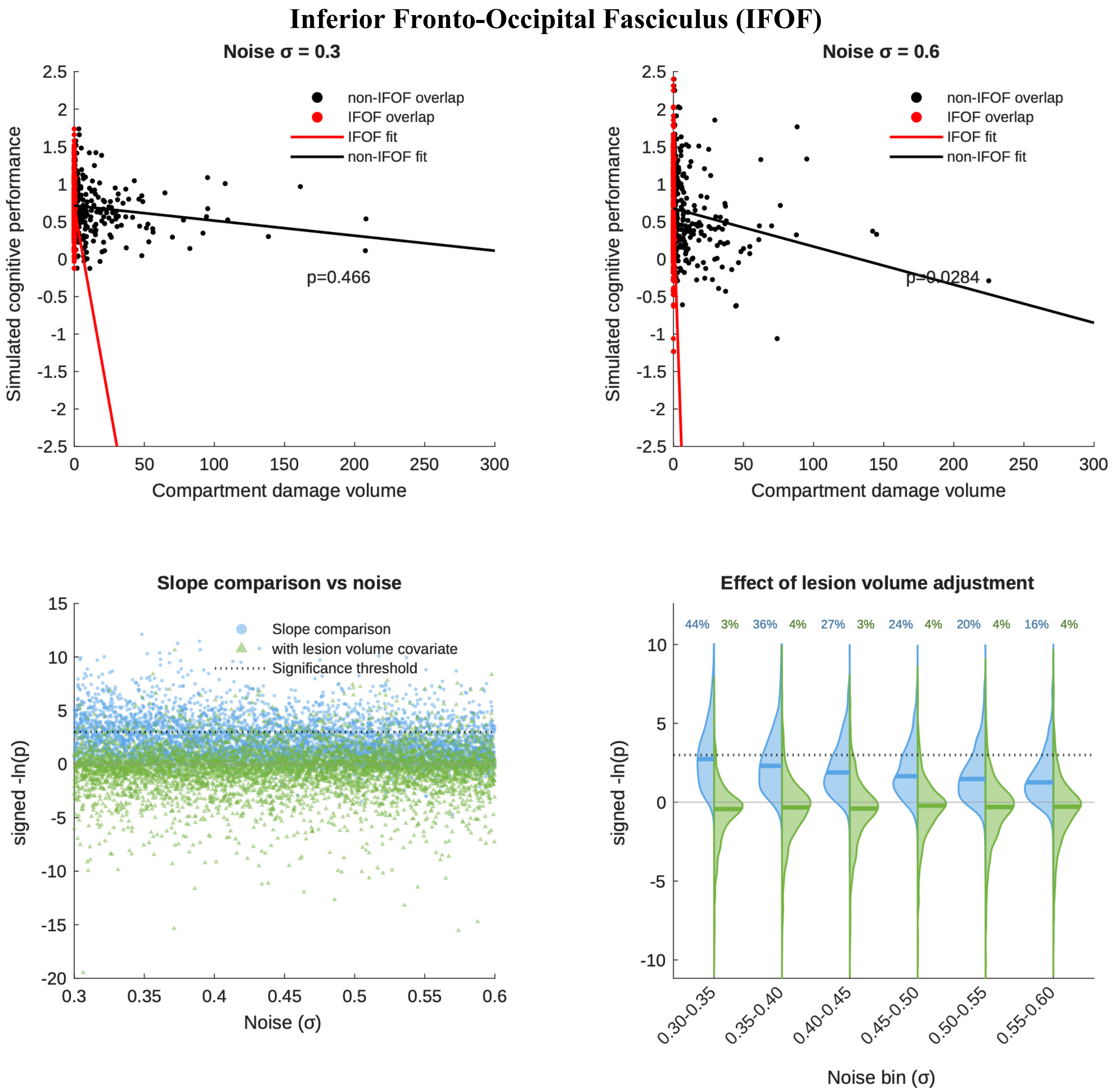


**Figure S7. Semi-synthetic simulations of RLDR with the Inferior Fronto-Occipital Fasciculus (IFOF) as the hypothesized substrate and lesion-size dependent deficit.** Example RLDR simulations of a purely lesion-size dependent deficit at two different noise levels (σ=0.3, top left; σ=0.6, top right), showing linear fits to IFOF (red) and non- IFOF (black) compartmental volumes, and the p value for the difference between the slopes. The plots below show the p values (expressed as their negative log) from 5000 Monte Carlo simulations across the full noise range, with (green) and without (blue) the addition of lesion volume as a covariate in the model, plotted individually (bottom left), and binned as violin plots (bottom right). Note that the p value distributions exhibit substantial mass surviving the critical threshold of $p<0.05$ (dotted line), quantified in the binned plots separately for volume-adjusted and unadjusted simulations, resulting in frequent spurious inference of critical IFOF involvement where there is none.

#### 2. True substrate associated semi-synthetic simulations

Semi-synthetic simulations with a true association between behaviour and the corresponding neural substrate map across various cortical networks and white matter tracts as described in the manuscript (see Figure 5).

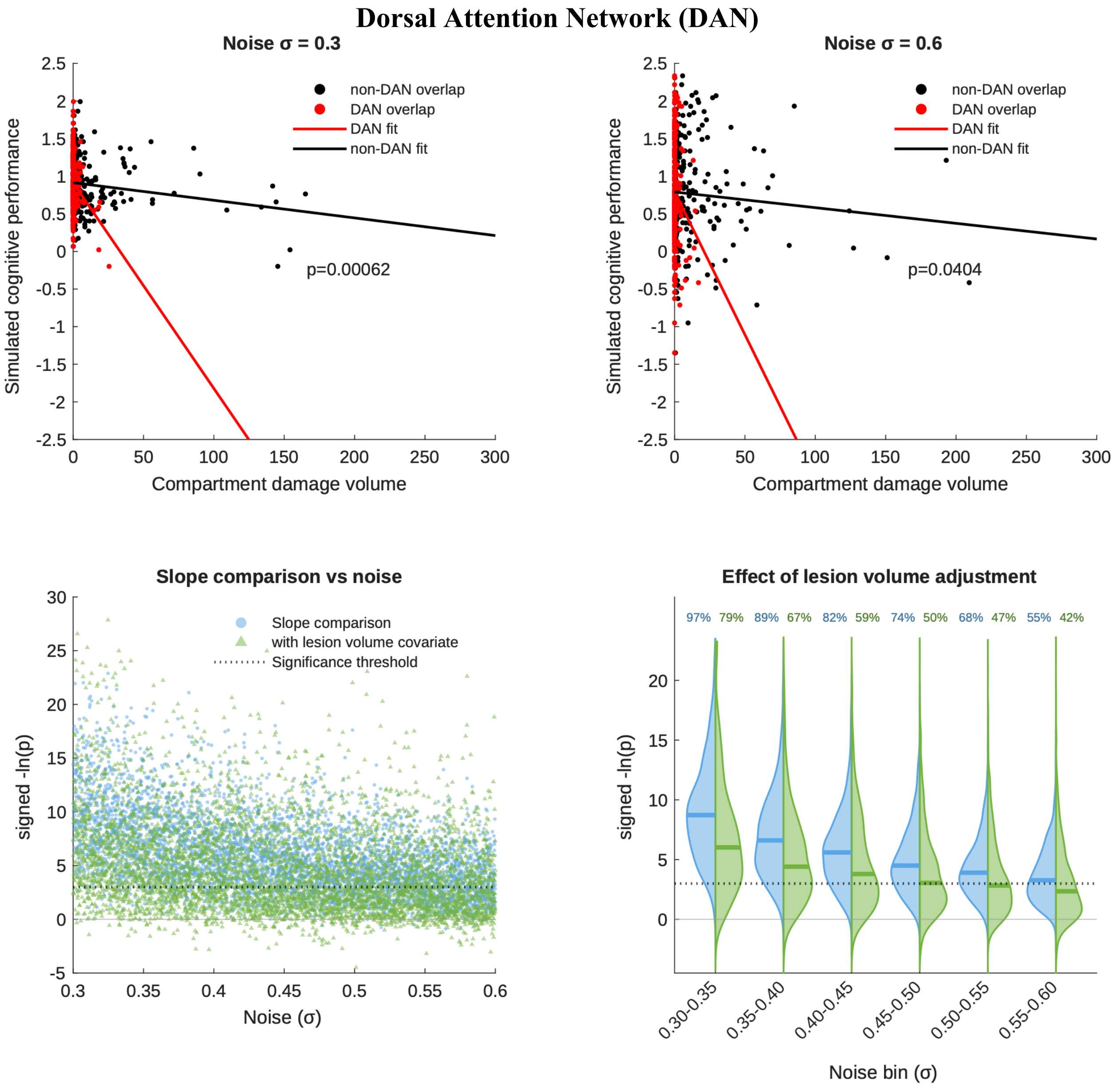


**Figure S8. Semi-synthetic simulations of RLDR with the Dorsal Attention Network (DAN) as the hypothesized and true substrate.** Example RLDR simulations of a DAN-dependent deficit at two different noise levels (σ=0.3, top left; σ=0.6, top right), showing linear fits to DAN (red) and non-DAN (black) compartmental volumes, and the p value for the difference between the slopes. The plots below show the p values (expressed as their negative log) from 5000 Monte Carlo simulations across the full noise range, with (green) and without (blue) the addition of lesion volume as a covariate in the model, plotted individually (bottom left), and binned as violin plots (bottom right). Note that the p value distributions exhibit substantial mass above the critical threshold of $p<0.05$ (dotted line), quantified in the binned plots separately for volume-adjusted and unadjusted simulations, resulting in poor sensitivity for detecting true dependence, especially with adjustment for lesion volume.

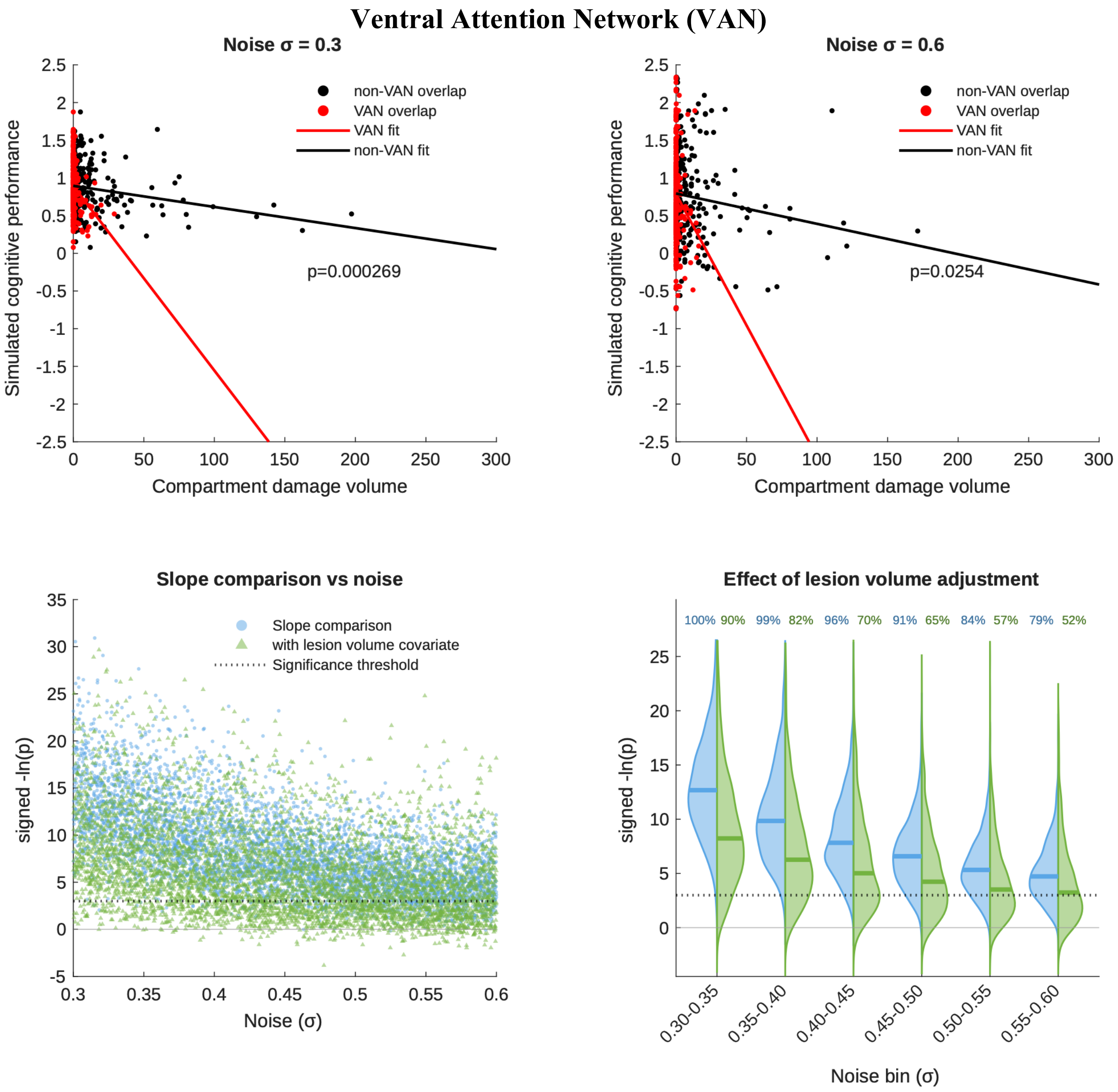


**Figure S9. Semi-synthetic simulations of RLDR with the Ventral Attention Network (VAN) as the hypothesized and true substrate.** Example RLDR simulations of a VAN-dependent deficit at two different noise levels (σ=0.3, top left; σ=0.6, top right), showing linear fits to VAN (red) and non-VAN (black) compartmental volumes, and the p value for the difference between the slopes. The plots below show the p values (expressed as their negative log) from 5000 Monte Carlo simulations across the full noise range, with (green) and without (blue) the addition of lesion volume as a covariate in the model, plotted individually (bottom left), and binned as violin plots (bottom right). Note that the p value distributions exhibit substantial mass above the critical threshold of $p<0.05$ (dotted line), quantified in the binned plots separately for volume-adjusted and unadjusted simulations, resulting in poor sensitivity for detecting true dependence, especially with adjustment for lesion volume.

### Sensorimotor Network (SM)

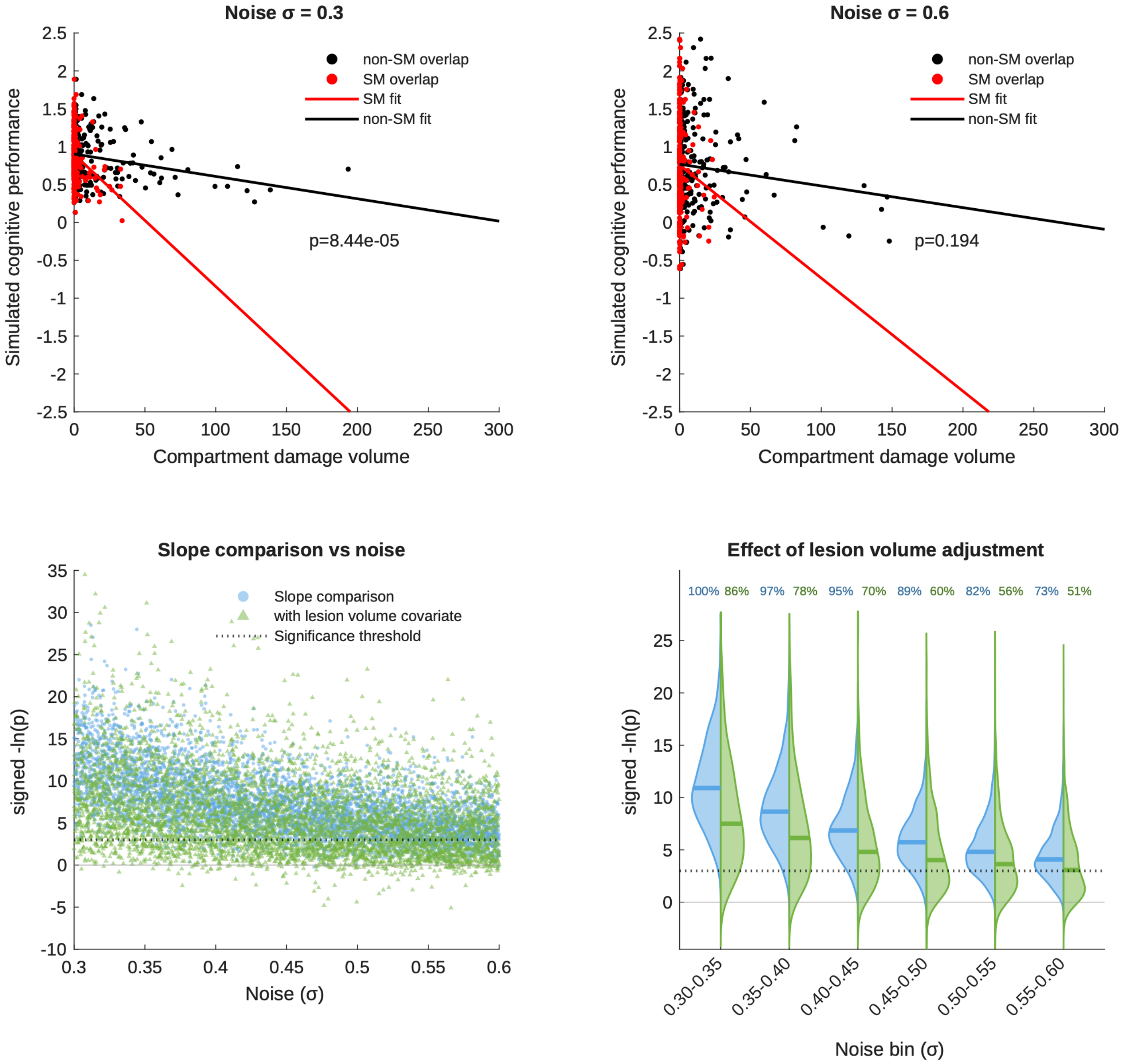


**Figure S10. Semi-synthetic simulations of RLDR with the Sensorimotor Network (SM) as the hypothesized and true substrate.** Example RLDR simulations of an SM-dependent deficit at two different noise levels (σ=0.3, top left; σ=0.6, top right), showing linear fits to SM (red) and non-SM (black) compartmental volumes, and the p value for the difference between the slopes. The plots below show the p values (expressed as their negative log) from 5000 Monte Carlo simulations across the full noise range, with (green) and without (blue) the addition of lesion volume as a covariate in the model, plotted individually (bottom left), and binned as violin plots (bottom right). Note that the p value distributions exhibit substantial mass above the critical threshold of $p<0.05$ (dotted line), quantified in the binned plots separately for volume-adjusted and unadjusted simulations, resulting in poor sensitivity for detecting true dependence, especially with adjustment for lesion volume.

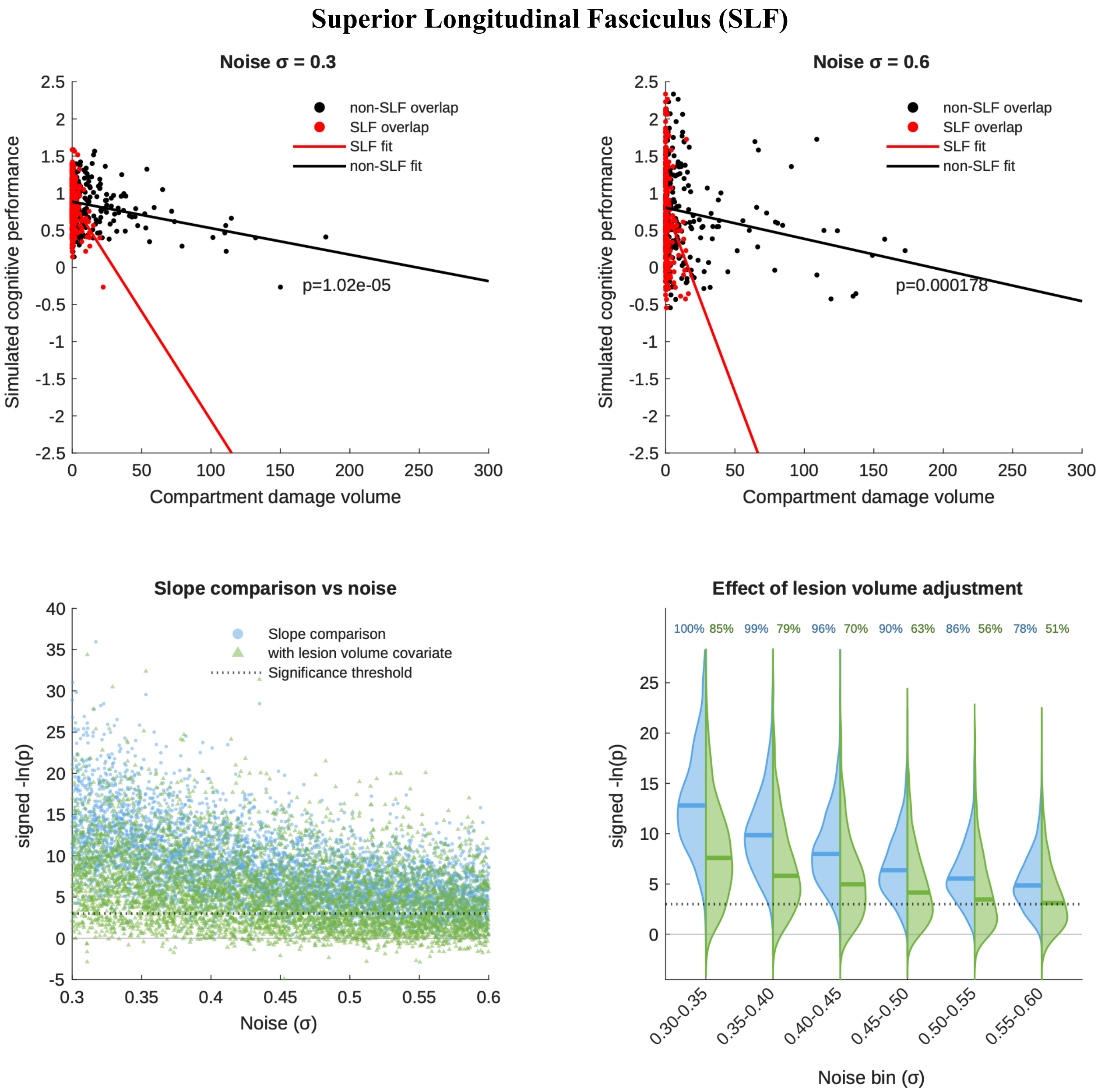


**Figure S11. Semi-synthetic simulations of RLDR with the Superior Longitudinal Fasciculus (SLF) as the hypothesized and true substrate.** Example RLDR simulations of a SLF-dependent deficit at two different noise levels (σ=0.3, top left; σ=0.6, top right), showing linear fits to SLF (red) and non-SLF (black) compartmental volumes, and the p value for the difference between the slopes. The plots below show the p values (expressed as their negative log) from 5000 Monte Carlo simulations across the full noise range, with (green) and without (blue) the addition of lesion volume as a covariate in the model, plotted individually (bottom left), and binned as violin plots (bottom right). Note that the p value distributions exhibit substantial mass above the critical threshold of $p<0.05$ (dotted line), quantified in the binned plots separately for volume-adjusted and unadjusted simulations, resulting in poor sensitivity for detecting true dependence, especially with adjustment for lesion volume.

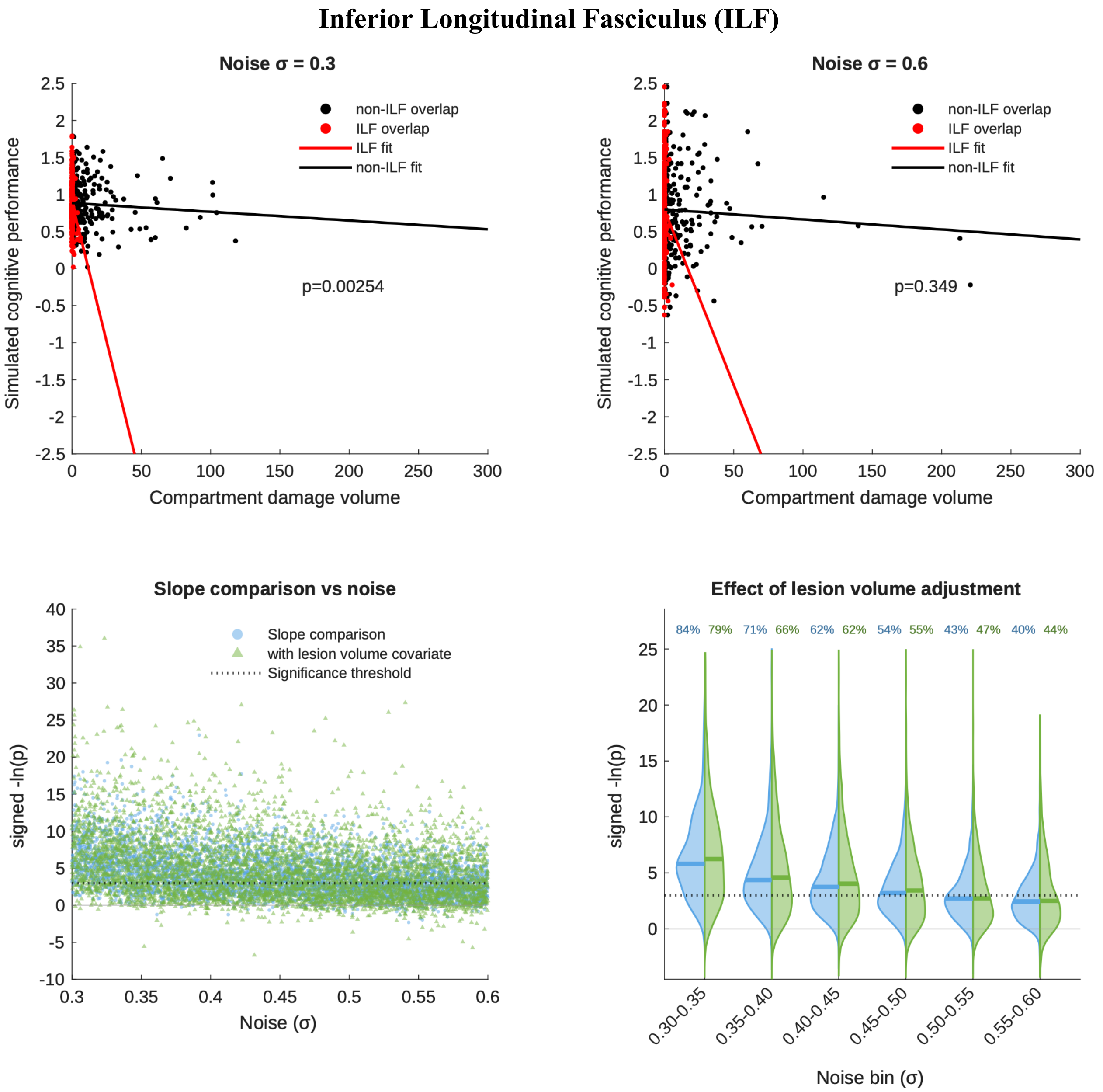


**Figure S12. Semi-synthetic simulations of RLDR with the Inferior Longitudinal Fasciculus (ILF) as the hypothesized and true substrate.** Example RLDR simulations of an ILF-dependent deficit at two different noise levels (σ=0.3, top left; σ=0.6, top right), showing linear fits to ILF (red) and non-ILF (black) compartmental volumes, and the p value for the difference between the slopes. The plots below show the p values (expressed as their negative log) from 5000 Monte Carlo simulations across the full noise range, with (green) and without (blue) the addition of lesion volume as a covariate in the model, plotted individually (bottom left), and binned as violin plots (bottom right). Note that the p value distributions exhibit substantial mass above the critical threshold of $p<0.05$ (dotted line), quantified in the binned plots separately for volume-adjusted and unadjusted simulations, resulting in poor sensitivity for detecting true dependence, especially with adjustment for lesion volume.

#### Arcuate Fasciculus (AF)

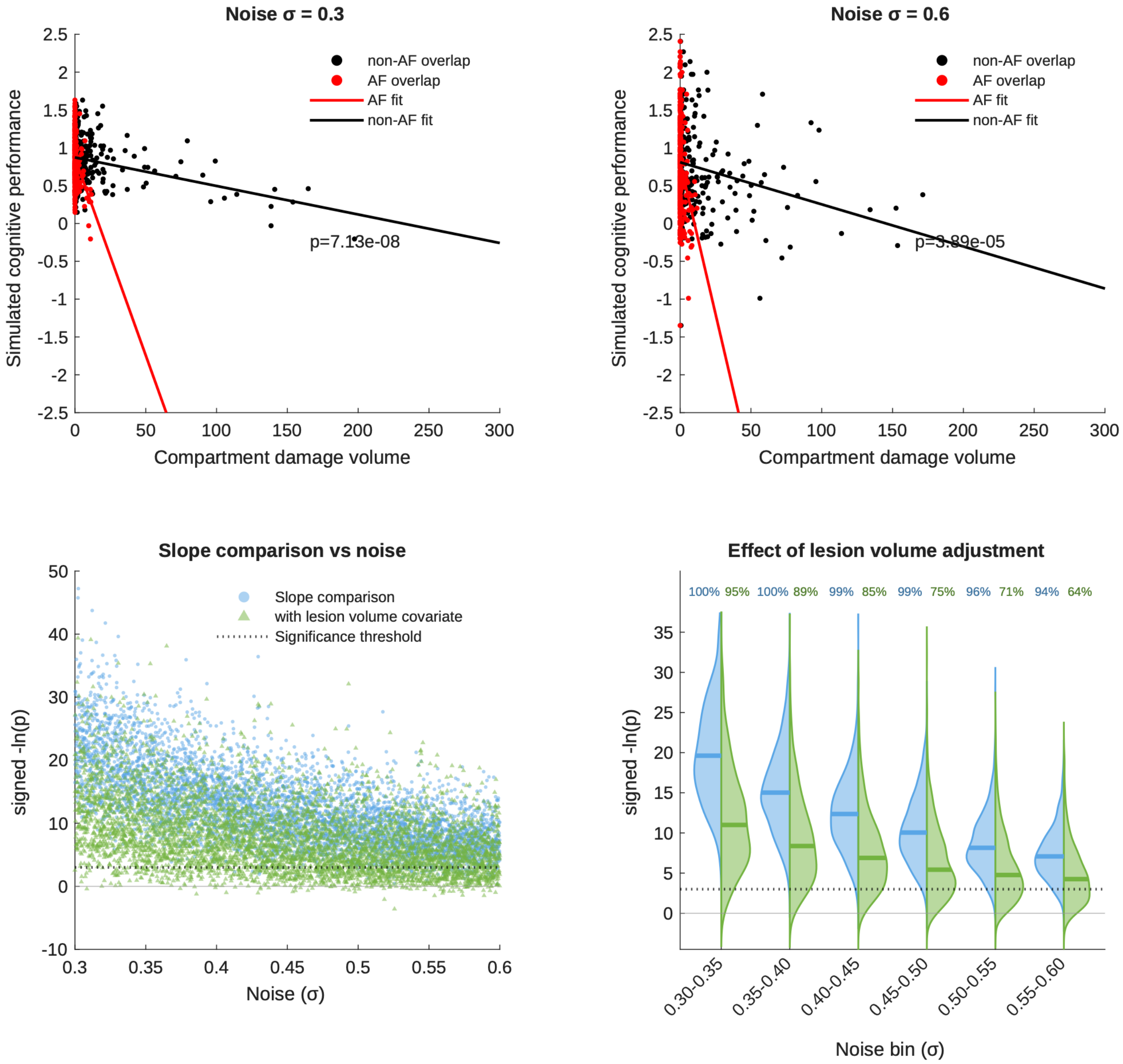


**Figure S13. Semi-synthetic simulations of RLDR with the Arcuate Fasciculus (AF) as the hypothesized and true substrate.** Example RLDR simulations of an AF-dependent deficit at two different noise levels (σ=0.3, top left; σ=0.6, top right), showing linear fits to AF (red) and non-AF (black) compartmental volumes, and the p value for the difference between the slopes. The plots below show the p values (expressed as their negative log) from 5000 Monte Carlo simulations across the full noise range, with (green) and without (blue) the addition of lesion volume as a covariate in the model, plotted individually (bottom left), and binned as violin plots (bottom right). Note that the p value distributions exhibit substantial mass above the critical threshold of $p<0.05$ (dotted line), quantified in the binned plots separately for volume-adjusted and unadjusted simulations, resulting in poor sensitivity for detecting true dependence, especially with adjustment for lesion volume.

**Inferior Fronto-Occipital Fasciculus (IFOF**

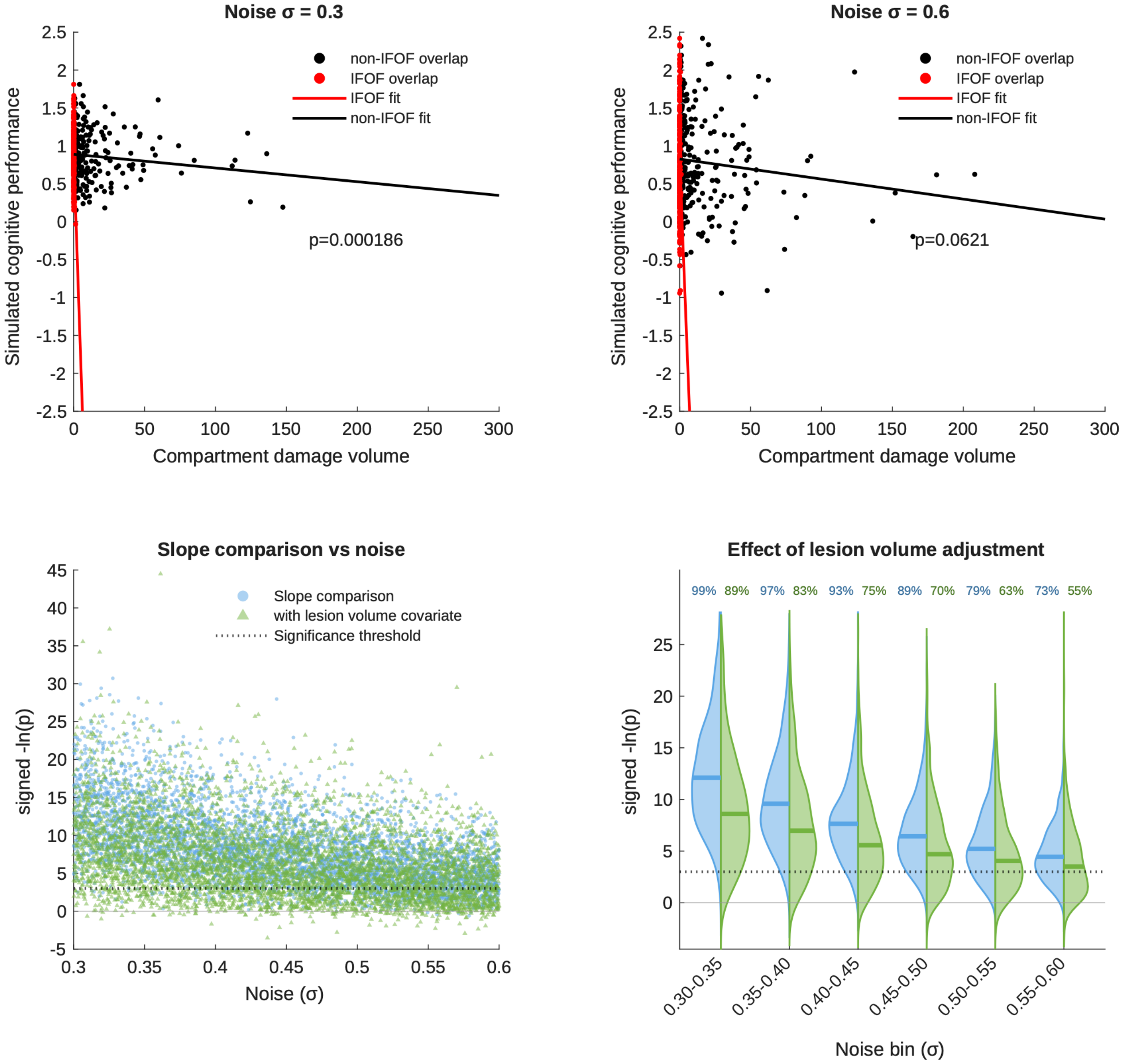


**Figure S14. Semi-synthetic simulations of RLDR with the Inferior Fronto-Occipital Fasciculus (IFOF) as the hypothesized and true substrate.** Example RLDR simulations of an Inferior Fronto-Occipital Fasciculus-dependent deficit at two different noise levels (σ=0.3, top left; σ=0.6, top right), showing linear fits to IFOF (red) and non-IFOF (black) compartmental volumes, and the p value for the difference between the slopes. The plots below show the p values (expressed as their negative log) from 5000 Monte Carlo simulations across the full noise range, with (green) and without (blue) the addition of lesion volume as a covariate in the model, plotted individually (bottom left), and binned as violin plots (bottom right). Note that the p value distributions exhibit substantial mass above the critical threshold of $p<0.05$ (dotted line), quantified in the binned plots separately for volume-adjusted and unadjusted simulations, resulting in poor sensitivity for detecting true dependence, especially with adjustment for lesion volume.
